# Pan-genome analysis highlights the role of structural variation in the evolution and environmental adaptation of *Asian honeybees*

**DOI:** 10.1101/2023.06.15.545041

**Authors:** Yancan Li, Jun Yao, Huiling Sang, Quangui Wang, Long Su, Xiaomeng Zhao, Zhenyu Xia, Feiran Wang, Kai Wang, Delong Lou, Guizhi Wang, Robert M. Waterhouse, Huihua Wang, Shudong Luo, Cheng Sun

**Affiliations:** State Key Laboratory of Resource Insects, Institute of Apicultural Research, Chinese Academy of Agricultural Sciences, Beijing, China; College of Life Sciences, Capital Normal University, Beijing, China; Western Research Institute, Chinese Academy of Agricultural Sciences, Changji, China; Institute of Plant Protection, Shandong Academy of Agricultural Sciences, Jinan, China; College of Animal Science, Shanxi Agricultural University, Shanxi, China; Shandong Provincial Animal Husbandry Station, Jinan, China; Department of Animal Science, Shandong Agricultural University, Taian, China; Department of Ecology and Evolution, University of Lausanne, and SIB Swiss Institute of Bioinformatics, 1015 Lausanne, Switzerland

## Abstract

The Asian honeybee, *Apis cerana*, is an ecologically and economically important pollinator. Mapping its genetic variation is key to understanding population-level health, histories, and potential capacities to respond to environmental changes. However, most efforts to date were focused on single nucleotide polymorphisms (SNPs) based on a single reference genome, thereby ignoring larger-scale genomic variation. We employed long-read sequencing technologies to generate a chromosome-scale reference genome for the ancestral group of *A. cerana*. Integrating this with 525 resequencing datasets, we constructed the first pan-genome of *A. cerana*, encompassing almost the entire gene content. We found that 31.32% of genes in the pan-genome were variably present across populations, providing a broad gene pool for environmental adaptation. We identified and characterized structural variations (SVs) and found that they were not closely linked with SNP distributions, however, the formation of SVs was closely associated with transposable elements. Furthermore, phylogenetic analysis using SVs revealed a novel *A. cerana* ecological group not recoverable from the SNP data. Performing environmental association analysis identified a total of 44 SVs likely to be associated with environmental adaptation. Verification and analysis of one of these, a 330 bp deletion in the *Atpalpha* gene, indicated that this SV may promote the cold adaptation of *A. cerana* by altering gene expression. Taken together, our study demonstrates the feasibility and utility of applying pan-genome approaches to map and explore genetic feature variations of honeybee populations, and in particular to examine the role of SVs in the evolution and environmental adaptation of *A. cerana*.

## Introduction

Honeybees are economically important insects that play an important role in honey production and agricultural pollination [1]. The Asian honeybee (*Apis cerana*) has the widest natural distribution among all Asian bee species, with habitats covering a wide range of environments: from tropical to cold temperate climates and from plains to mountains [2]. As a result of long-term natural selection, *A. cerana* has adapted well to local environments, and the wide distribution pattern suggests that *A. cerana* could be used to study the molecular basis underlying its adaptation to different environments [3]. In addition, compared to western honeybees (e.g., *Apis mellifera*), *A. cerana* is less affected by the confounding signals produced by genetic mixing caused by artificial interventions (e.g., artificial breeding and domestication) and has maintained a semiwild feature [2-4], which further makes it a good model organism for studying the genetic mechanisms underlying adaptation. Furthermore, the population of *A. cerana* has declined and is declining [5-8], so understanding how *A. cerana* populations adapt to their external environmental conditions and elucidating the underlying mechanisms could greatly inform conservation efforts.

Several large-scale whole-genome resequencing projects have been performed for *A. cerana*, which not only revealed its population genetic structure but also uncovered some genes linked to its adaptations [3, 9, 10]. However, previous analyses mainly used a single reference genome and mainly focused on small-sized genetic variations, such as single nucleotide polymorphisms (SNPs) and small insertions/deletions (Indels). Single reference genomes, especially those generated by short-read sequencing, often fail to detect complete genomic variation and thus could miss important variation information [11, 12]. For example, a pan-genome study of African humans found that nearly 10% of the sequences were missing from the reference genome [13]. In addition, SNPs or Indels cannot represent the complete genetic variation repertoire of one species, and other larger-sized genetic variations, such as presence/absence variations (PAVs) and structural variations (SVs), could also play important roles in the adaptation of organisms (including insects) to diverse environments [14-16]. Therefore, to delineate a comprehensive repertoire of variants and identify all possible causal variations accounting for adaptation, we need to construct a pan-genome (the collection of all DNA sequences for a species) for *A. cerana* and genotype larger-sized genetic variations through population genomics studies.

In this study, we first constructed and characterized the pan-genome of *A. cerana* based on a chromosome-level reference genome for its ancestral group and 525 whole-genome resequencing datasets. Then, we investigated the presence/absence variation (PAV) patterns and genotyped structural variations (SVs) across *A. cerana* populations. Finally, we used these mapped variants to explore the potential role of SVs in the environmental adaptation of *A. cerana*. These resources achieve a milestone in the application of genomics technologies to characterise and explore the complete pan-genome content and how it varies across populations of this important pollinator.

## Results

### Chromosome-scale reference genomes for several groups of Asian honeybees

The population structure of *A. cerana* is composed of one central ancestral group and multiple peripheral groups that radiated from it [3, 17]. In this study, we generated a chromosome-scale reference genome sequence for the ancestral group of *A. cerana* (named Hubei to indicate samples were collected from the Hubei Province in China) by applying PacBio high-fidelity (HiFi) sequencing and high-throughput chromosome conformation capture (Hi-C) techniques **(Fig. 1 A; Supplementary Tab. 2)**. The resultant genome assembly comprised 16 chromosome-scale scaffolds (**Fig. 1 B**), with over 99% of contigs placed on these scaffolds (**Supplementary Tab. 3**). Furthermore, 83.02% of the scaffold sequences from the ACSNU-2.0 *A. cerana* genome were reflected in this chromosome (**Fig. 1 C**; identity > 0.75). The final genome size and scaffold N50 were 217.7 Mb and 4.15 Mb, respectively (**Fig. 1 E**; **Supplementary Tab. 2**). BUSCO analysis [18] indicated that the genome assembly had a high completeness score (98.5%) and was substantially more complete than the frequently used ACSNU-2.0 reference genome [19] (**Fig. 1 E**; **Supplementary Tab. 4**), which has been widely used in previous population genomics analyses [3, 9, 20, 21]. The newly generated reference genome was annotated by integrating multiple lines of evidence (**see Methods**), and a total of 11,362 protein-coding genes were annotated, of which 10,073 were assigned at least one Gene Ontology (GO) term or protein domain (**Supplementary Tab. 4; Supplementary Table 8**).

**Fig. 1:**
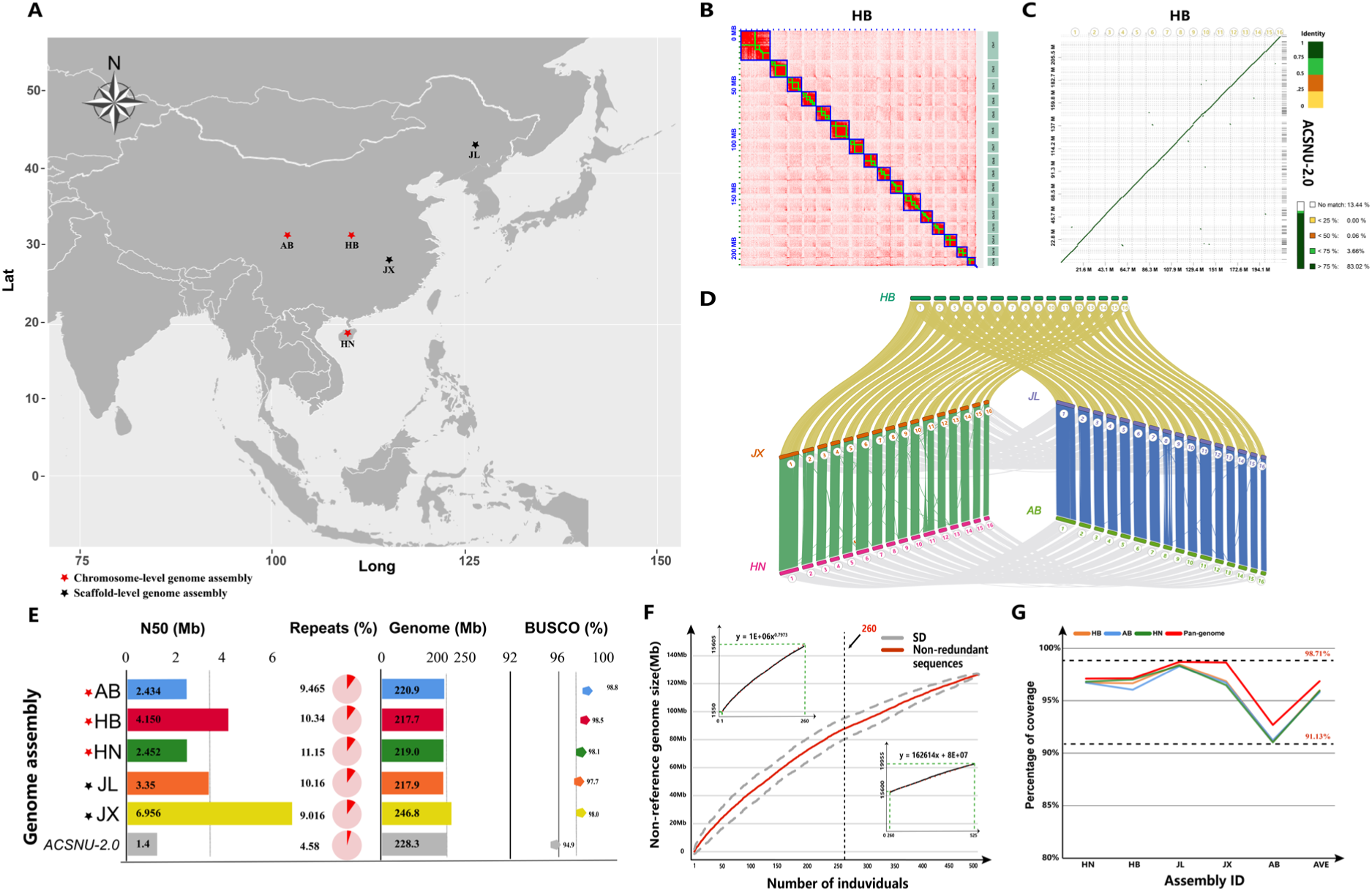
Reference genomes and pan-genome generated for Asian honeybees. **A.** The geographic distribution of *A. cerana* samples that were used for reference genome construction (AB, HB, HN, JL, and JX, which are shorthand names for Aba/Sichuan, Hubei, Hainan, Jilin, and Jiangxi, respectively). Red stars represent chromosome-level genome assemblies, while black stars indicate scaffold-level genome assemblies. **B**. The Hi-C heatmap for the chromosome-level reference genome of the *A. cerana* ancestral group (named Hubei). **C.** Synteny between the ACSN-2.0 genome and the Hubei genome. Four thresholds were set according to different similarities, which were 75%-100% (dark green), 50%-75% (green), 25%-50% (orange), and 0%-25% (yellow), with white representing the unmatched area. The analysis revealed that 82.87% of the ACSN-2.0 genome was identified in the chromosomes of the newly assembled genome. **D**. Chromosome collinearity across the five *A. cerana* genomes assembled in this study (HB, JX, HN, JL and AB). Chromosomes were numbered from Chr1 to Chr16 based on their synteny with the *Apis mellifera* genome (Amel_HAv3.1). **E.** Comparison of newly generated reference genomes with the frequently used reference genome ACSNU-2.0. N50 (Mb): scaffold N50 length in Mb; Repeats (%): the percentage of the reference genome that was recognized as repetitive sequences; Genome (Mb): genome size in Mb; BUSCO (%): genome completeness in % based on BUSCO analysis with the Hymenoptera dataset. **F.** The construction of the *A. cerana* pan-genome. The length of nonreference sequences increased when new *A. cerana* individuals were added. If the total number of individuals is less than 260, the length of nonreference sequences has an exponential growth, and if the number of individuals is larger than 260, it has a linear growth. As shown, red lines are nonredundant nonreference sequences, and SD is the standard deviation. **G**. Evaluation of the pan-genome assembly by mapping long reads of five individuals onto the reference genome and the pan-genome, respectively. From left to right are Hainan, Hubei, Jilin, Jiangxi, Aba, and average.

We also generated genome assemblies using long-read sequencing for four peripheral groups of *A. cerana* (namely, Aba, Jilin, Jiangxi, and Hainan) with BUSCO completeness ranging from 97.70% to 98.80% (**Fig. 1 D, E; Supplementary Tab. 2**), among them, the Aba and Hainan assemblies were chromosomal-level collections generated with the assistance of Hi-C data (**Supplementary Fig. 1 A, B**). Repetitive sequences (including transposable elements) were annotated in all five genome assemblies, and the repeat content ranged from 9.02% to 11.15% (**Fig. 1 E; Supplementary Fig. 2; Supplementary Tab. 5**). All five genome assemblies showed higher genome continuity, gene completeness, and repeat content than the frequently used *A. cerana* reference genome ACSNU-2.0 (**Fig. 1. C, E**). Taken together, we generated several high-quality genome assemblies for multiple groups of *A. cerana*, which represent valuable genomic resources for future studies.

### The features of the Asian honeybee pan-genome

A reference-guided assembly approach [14, 22] was used to construct the *A. cerana* pan-genome. In brief, whole-genome shotgun read (WGS) datasets of 525 *A. cerana* individuals were aligned to the reference genome of the ancestral group of *A. cerana* (Hubei), and unmapped reads were subjected to *de novo* assembly, followed by filtering contamination and redundancy (**for a detailed pipeline, see Supplementary Fig. 3 and Supplementary Tab. 6**). The size of the pan-genome increased rapidly at the beginning and gradually slowed as the number of individuals increased, with an inflection point of approximately 260 individuals, above which the pan-genome size increased linearly at a rate of approximately 0.163 Mb per individual (**Fig. 1 D**). After iterative addition of the 525 *A. cerana* individuals, 127.6 Mb of non-redundant, non-reference sequences were obtained, resulting in a final *A. cerana* pan-genome of 345.2 Mb in length, with 217.7 Mb of reference and 127.6 Mb of nonreference sequences (**Fig. 1 D**). A total of 16,587 protein-coding genes were annotated in this pan-genome. The completeness of the *A. cerana* pan-genome was evaluated using three chromosome-scale *A. cerana* reference genomes employing the read-back mapping method (**see Methods**), and the pan-genome was improved by 0.96% on average (the average mapping rate of the three reference genomes was 95.90%, and the average mapping rate of the *A. cerana* pan-genome was 96.86%) (**Fig. 1 E and Supplementary Tab. 7**). That is, the *A. cerana* pan-genome we generated in this study is more representative than any single chromosome-level reference genome.

After obtaining the annotated genes of the pan-genome (pan-genes), gene presence-absence variation (PAV) patterns for each *A. cerana* individual were estimated by mapping their WGS data to the 16,587 pan-genes (**see Methods**). To ensure accuracy, a total of 502 high-quality resequencing datasets were selected for this analysis (**Supplementary Tab. 1**). The total gene set increased when additional *A. cerana* individuals were added but gradually approached a plateau when n = 351 (99% of *A. cerana* pan-genes, **Fig. 2 A**), indicating that we obtained a “closed” set of pan-genes using the currently available dataset. The PAV matrix showed a high genotyping accuracy (99.12% for true presence and 90.71% for true absence, **Supplementary Fig. 4**), and we categorized the 16,587 genes in the pan-genome according to their frequency of occurrence using previous standards [14, 16]. A total of 8,544 (51.51%) genes were shared by all *A. cerana* individuals, which were classified as core genes; 2,815 (16.97%) genes were categorized as softcore genes occurring in 377– 502 individuals (75–100%); 2,381 (14.35%) genes were named shell genes present in 15–377 individuals (3–75%); 1,199 (7.23%) genes were classified as cloud genes present in 5-15 individuals (1%-3%); leaving 1,648 single genes, which were found in fewer than 5 individuals (less than 1%) (**Supplementary Fig. 5 A-D**). Based on genome annotation results, we found that 1) genes located on chromosomes had more exons and longer transcripts; 2) all core genes were located on 16 chromosomes; and 3) while most of the softcore genes were located in the reference genome sequence, most shell genes were present in nonreference sequences. Cloud genes and single genes were located entirely in nonreference sequences, which only occurred in some of the *A. cerana* populations (**Supplementary Tab. 8; Supplementary Fig. 5 A, E-F**).

**Fig 2:**
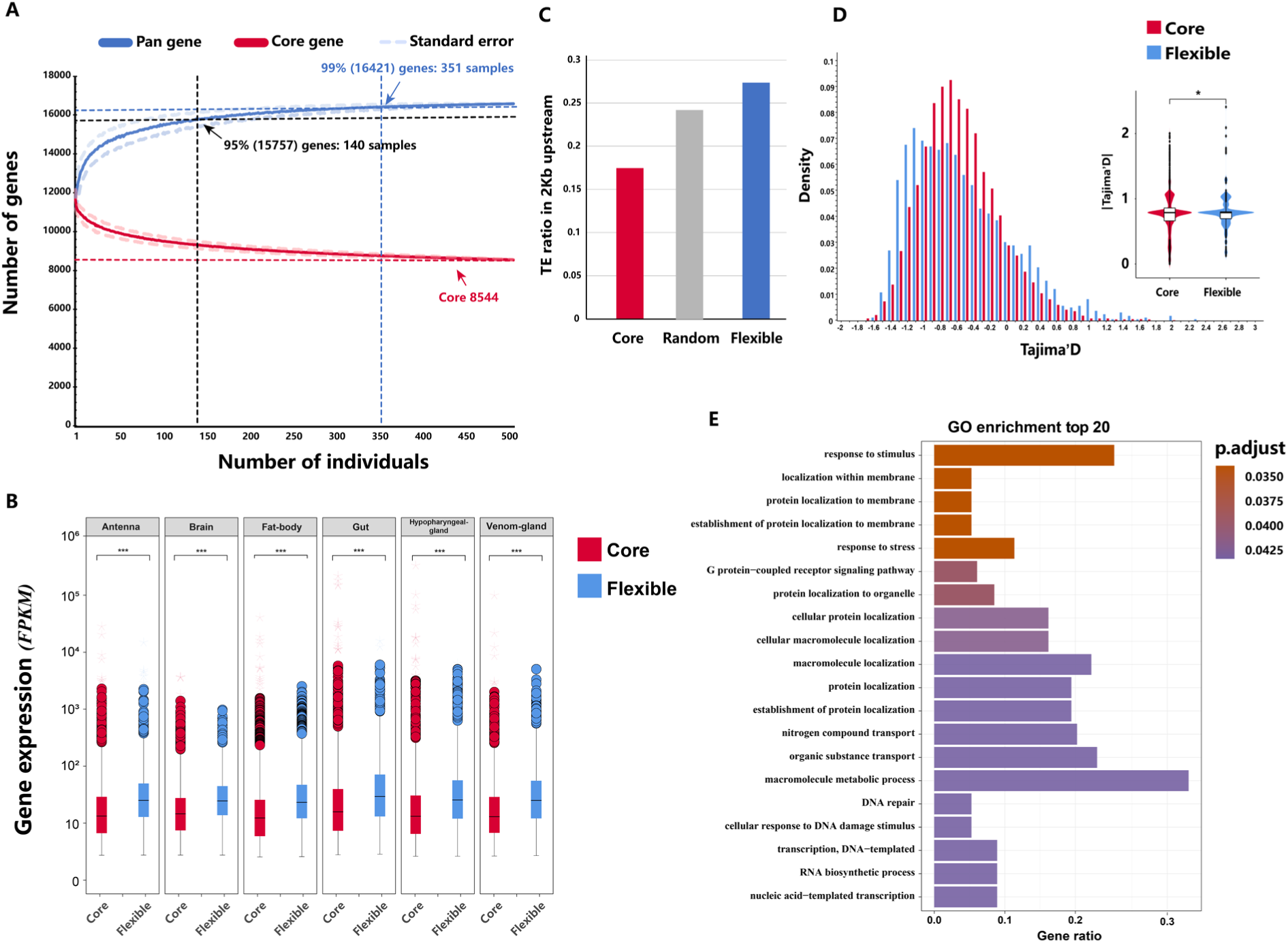
Features of the *A. cerana* pan-genome. **A.** Visualization of the number of pan (blue) and core (red) genes obtained by using different numbers of sequenced *A. cerana* individuals. The red dashed line represents the number of 99% of all pan-genome genes, and the black dashed line represents the number of genes at 95% of all pan-genome genes. **B.** Box plots show the expression levels of core and flexible genes in different tissues (including antenna, brain, fat body, gut, hypopharyngeal gland, venom gland), with circles representing outliers and stars representing extremes. Transcriptome data were downloaded from the NCBI SRA database (see Methods). The comparison of gene expression and Tajima’s D between core and flexible genes was carried out using the Wilcoxon test (*P < 0.05, **P < 0.01, ***P < 0.001). **C.** Ratio of transposable element (TE) insertion frequencies in the 2 kb upstream of core and variable genes. To evaluate bias in the actual TE distribution, we created 1000 randomly shuffled TE sets of the reference genome. **D.** The probability density plot of Tajima’s D in the sliding window shows the Tajima’s D values for core and flexible genes as well as the distribution of their absolute values. The selected genomic regions were identified by SNP data, and the figure shows that the flexible genes are left-skewed relative to the core genes, indicating that flexible genes are under greater positive selection. **E.** Gene Ontology (GO) enrichment analysis of flexible genes (only biological process (BP) entries are shown).

For the convenience of subsequent analysis, we collectively refer to softcore and shell genes as “flexible” genes. Likewise, we refer to “cloud genes” and “single genes” as “specific” genes. Based on the available *A. cerana* transcriptome data (**see Methods**), we found that flexible genes were expressed at higher levels than core genes, and they were more likely to have TE insertions (especially Helitron transposons) in their nearby regions (**Fig. 2 B-C; Supplementary Fig. 5 G-H**). Based on results from a genome-wide scan of selective regions using SNP data, we found that more flexible genes than core genes were under positive selective pressure (**Fig. 2 D**). Gene Ontology (GO) analysis showed that core genes were enriched in “house-keeping” biological processes, while the flexible and specific genes were enriched in biological processes such as “response to stress” and “response to stimulus”, indicating that they are likely involved in *A. cerana* adaptation (**Fig. 2 E; Supplementary Fig. 6**). Taken together, we have generated a pan-genome for *A. cerana* in which the protein-coding gene repertoire should be complete. Our results also indicated that gene loss/gain is common across *A. cerana* individuals, and the evolution of flexible genes is faster than that of core genes, which might be related to the adaptation of *A. cerana* to diverse habitats.

### Structural variants are closely associated with transposable elements

Whole-genome shotgun reads of 525 *A. cerana* individuals were mapped back to the ancestral group of the *A. cerana* reference genome (Hubei, HB) to identify SVs. To ensure the accuracy and reliability of the results, a combination of multiple software programs was used, and strict filtering criteria were applied (**see Methods**). The results showed that as the number of *A. cerana* individuals increased, the growth of the nonredundant set of SVs slowed down (**Supplementary Fig. 7 A**), which is similar to the process of pan-genome sequence construction, suggesting that most SVs should have been detected. A total of 19,955 nonredundant SVs were identified, with each *A. cerana* sample yielding an average of 1,550 SVs (**Fig. 3 A)**. The most common type of SV was deletion (DEL), and the majority of genotyped SVs were within 500 bp in length (90.64%, n = 17,584) (**Fig. 3 B; Supplementary Fig. 7 B**).

**Fig. 3:**
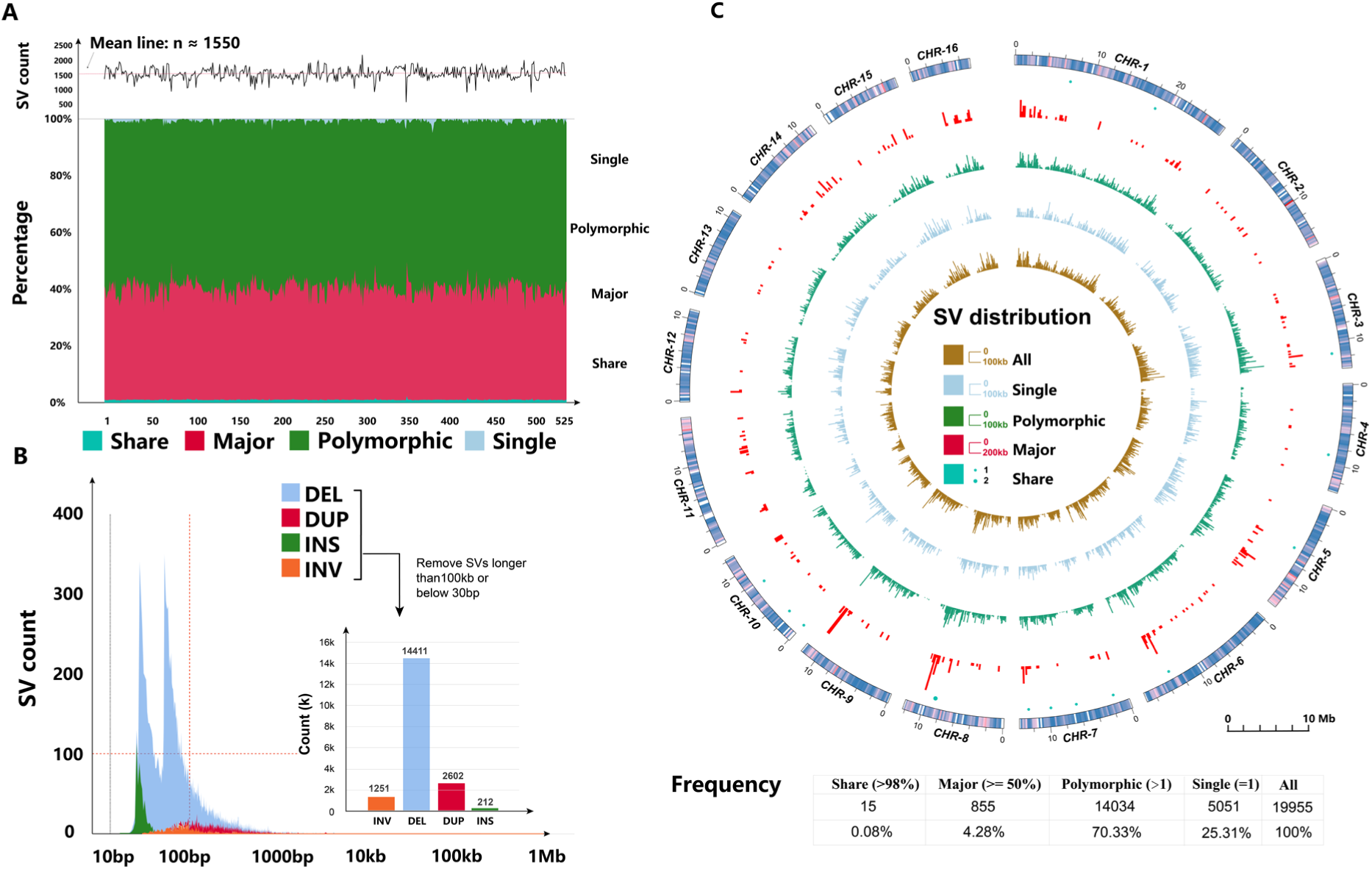
The number, length, and distribution of *A. cerana* SVs. **A.** The number of SVs contributed by each *A. cerana* individual in each of the following SV categories: shared SV (identified in all samples), major SV (identified in ≥50% of samples), polymorphic SV (identified in > 1 sample), and singleton SV (identified in only one sample). **B.** The length distribution of SVs for each SV type (DEL, deletion; DUP, duplication; INS, insertion; INV, inversion), as well as length distribution after filtering (SV length between 30 bp and 100 kbp were retained). Here, we did not count translocation (BND). **C.** The distribution of each SV category along *A. cerana* chromosomes, and the outermost layer shows the corresponding gene distribution on the same chromosome.

Based on previous criteria [23, 24], SVs were classified into four categories: shared (identified in ≥ 98% of samples), major (identified in ≥ 50% of samples), polymorphic (identified in > 1 sample), and singleton (identified in only one sample) (**Fig. 3 C**), and over half of the SVs were polymorphic (n = 14,034**; Supplementary Tab. 9; Supplementary Fig. 7 C-D**). Long-read sequencing reads were used to validate the accuracy of our genotyped SVs **(see Methods)**, and approximately 75% of the shared SVs were confirmed **(Supplementary Tab. 10)**. That is, although there was bias and misalignment during SV detection [25, 26], our genotyping results demonstrated decent accuracy.

The distribution of SVs in the *A. cerana* reference genome was analysed, and we found that while 78.29% of the identified SVs were located in intergenic regions (**Fig. 4 A; Supplementary Tab. 10**), 21.7% of SVs were distributed in genic regions (of which 84.29% were intronic regions), with only a small fraction of SVs located in exons **(Fig. 4 A**). Genomic regions that are under selection were identified based on SNP data, and we found that 7.97% of SVs were located in such regions (**Fig. 4 A; Supplementary Tab. 9**). Comparing SV and repeat genomic distributions, we found that 8.52% of SV breakpoints overlapped with repetitive elements, with DNA transposons (3.74%) and simple repeats (2.05%) being the dominant types (**Fig. 4 B**).

**Fig. 4:**
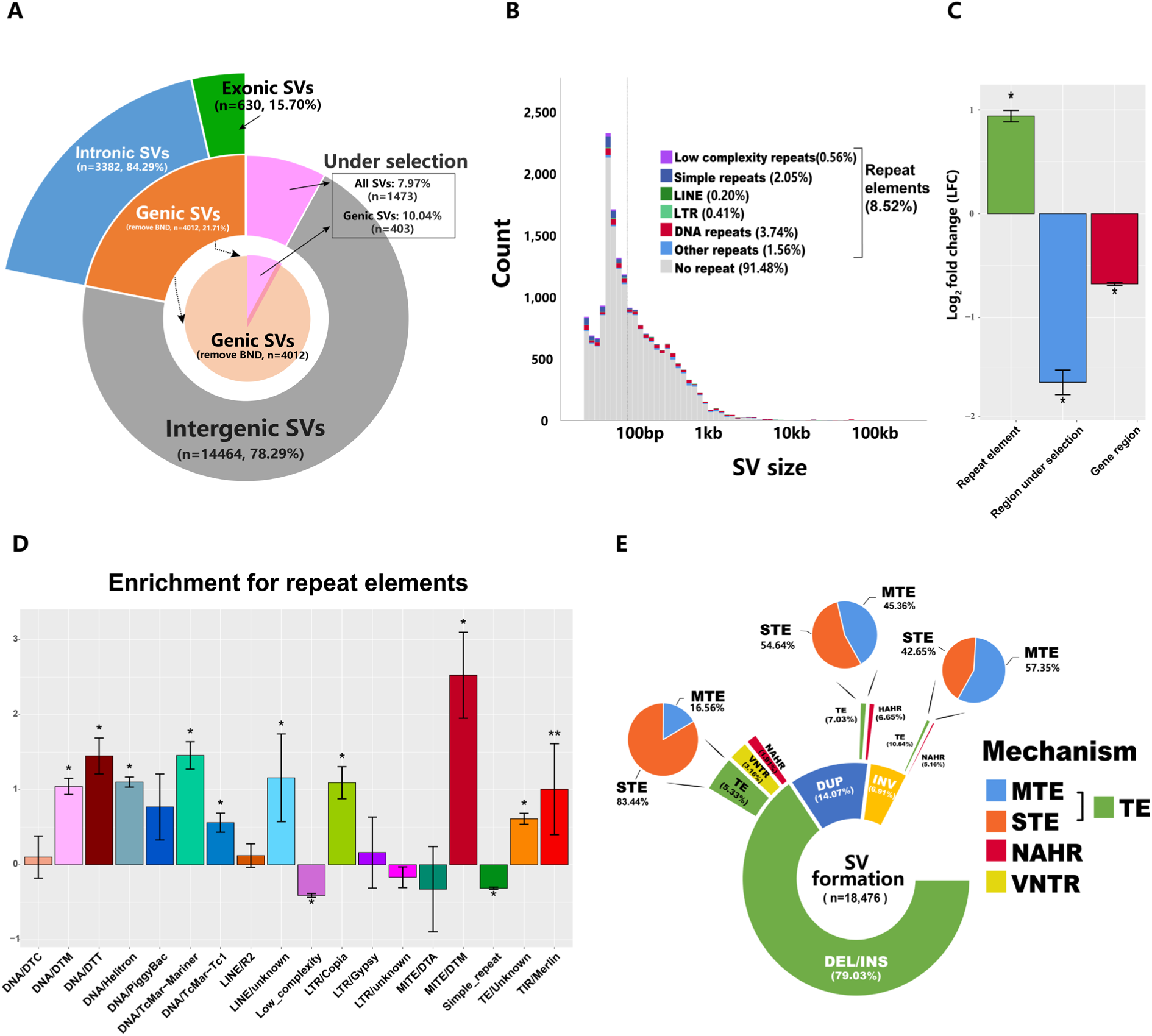
The distribution of SVs relative to genes, regions under selection, and repeats. **A.** The overlap of SVs with nearby genic regions and regions under selection. Regions under selection were obtained by calculating SNP data near those SVs. Note that translocations (BND) are excluded from this analysis. **B**. Bar charts show the overlap of SVs with repetitive sequences of different lengths. Repetitive sequences were further classified into low complexity repeats, simple repeats, long terminal repeat retrotransposons (LTR), non-LTR retrotransposons (LINE), DNA transposons, and other repeats. **C.** Random background obtained by 1,000 data simulations was compared to the true value of repetitive sequences, regions under selection and genic regions that overlapped with SVs, with values of the y-axis representing the log2-fold change of enrichment (LFC). Asterisks indicate significant enrichment with Bonferroni corrected p values < 0.05. **D.** Log2-fold change in enrichment analysis for repeat elements overlapping with the flanking sequence of the SV breakpoint (+/-100 bp). **E.** Inference of the mechanism of SV formation. SV types include deletions/insertions (DEL/INS), duplications (DUP), and inversions (INV). The SV formation mechanisms can be classified into four categories: nonallelic homologous recombination (NAHR), variable number tandem repeats (VNTR), single transposable element (STE), and multiple transposable element (MTE).

To detect biases in SV genomic distributions, we constructed a random background and found that SVs were significantly enriched in the repetitive regions (in terms of fold change; FC = 0.9, p value < 0.05) (**Fig. 4 C**). Examining the repetitive regions in detail, SVs preferred DNA transposons (DNA-TE), and the most enriched transposon was MITE/DTM (FC = 2.5, p value < 0.05; **Fig. 4 D**); there was a depletion of SVs in tandem repeat regions (FC < 0, low complexity and simple repeats) (**Fig. 4 D**). As reported, SVs occurring in functional sequences are usually deleterious and can disrupt gene function with phenotypic effects, so SVs located in genic regions should be eliminated during evolution [26-28]. As expected, compared to the repeat regions, the genic regions and genomic regions under selection exhibited a significant depletion of SVs (**Fig. 4 C; Supplementary Tab. 9**).

We employed a simplified pipeline as used in previous studies [23, 24, 29, 30] to infer the mechanism of SV formation based on breakpoints and their flanking sequence profiles (**Supplementary Fig. 8**). For all SV events (< 100 kbp, n = 18,476, except translocations (BND)), transposable elements (TE) were the dominant formation mechanism (5.33% of INS/DEL; 7.03% of DUP; 10.64% of INV); insertions and deletions were mainly mediated by a single transposon (83.44%), while duplications and inversions were mostly mediated by multiple transposons. In addition, variable number tandem repeat (VNTR) and nonallelic homologous recombination (NAHR) are also important mechanisms of SV formation: VNTR was mainly found in insertion/deletion SVs, while NAHR could promote the formation of duplications and inversions (**Fig. 4 D; Supplementary Tab. 9**).

As reported in several other species, the nucleotide diversity of SNPs and chromosome size are generally inversely related due to uncoupling effects [31-33]. We found that the general diversity of SVs in the *A. cerana* population may be driven by linked selection because a weak but significant negative correlation was detected between SV diversity (π) and chromosome size (**Supplementary Fig. 7 E;** R = -0.5; p = 0.052).

### SVs provide insights into the population structure of *A. cerana*

It has been shown in some species that SVs could provide insights into population structure [14, 34-36]. In this study, we reconstructed the population structure of *A. cerana* based on SVs and SNPs and compared the two results. First, the WGS data of 525 *A. cerana* samples were mapped to the reference genome generated in this study (Hubei). After quality control and pruning, ∼0.96 million SNPs were used to infer population structure using ADMIXTURE [37]. At the optimal K value of 9, nine distinct populations were obtained, with one central ancestral group (referred to as Central), comprising two components and several peripheral groups, namely Aba (AB), Bomi (BM), Central (CT), Hainan (HN), Northeast (NE), Qinghai (QH), Taiwan (TW), and a subpopulation called Malay (ML) (**Fig. 5 A and Supplementary Fig. 9 A-C**). This population structure is consistent with previous results using SNP data but a different reference genome [3, 17]. Then, we investigated the population structure based on SVs, which exhibited a finer population structure with an optimal K value of 10 (**Fig. 5 B; Supplementary Fig. 10 A**). Compared with results based on SNP data, a novel population of southern Yunnan (DN) was resolved at the transition from the CT to ML group **(Fig. 5 A-D; Supplementary Fig. 10 B-H**). The existence of the DN group is supported by a recent analysis based on morphological characters [38], indicating that SVs provide insights into the population structure of *A. cerana* beyond those obtainable from SNP data.

**Fig. 5:**
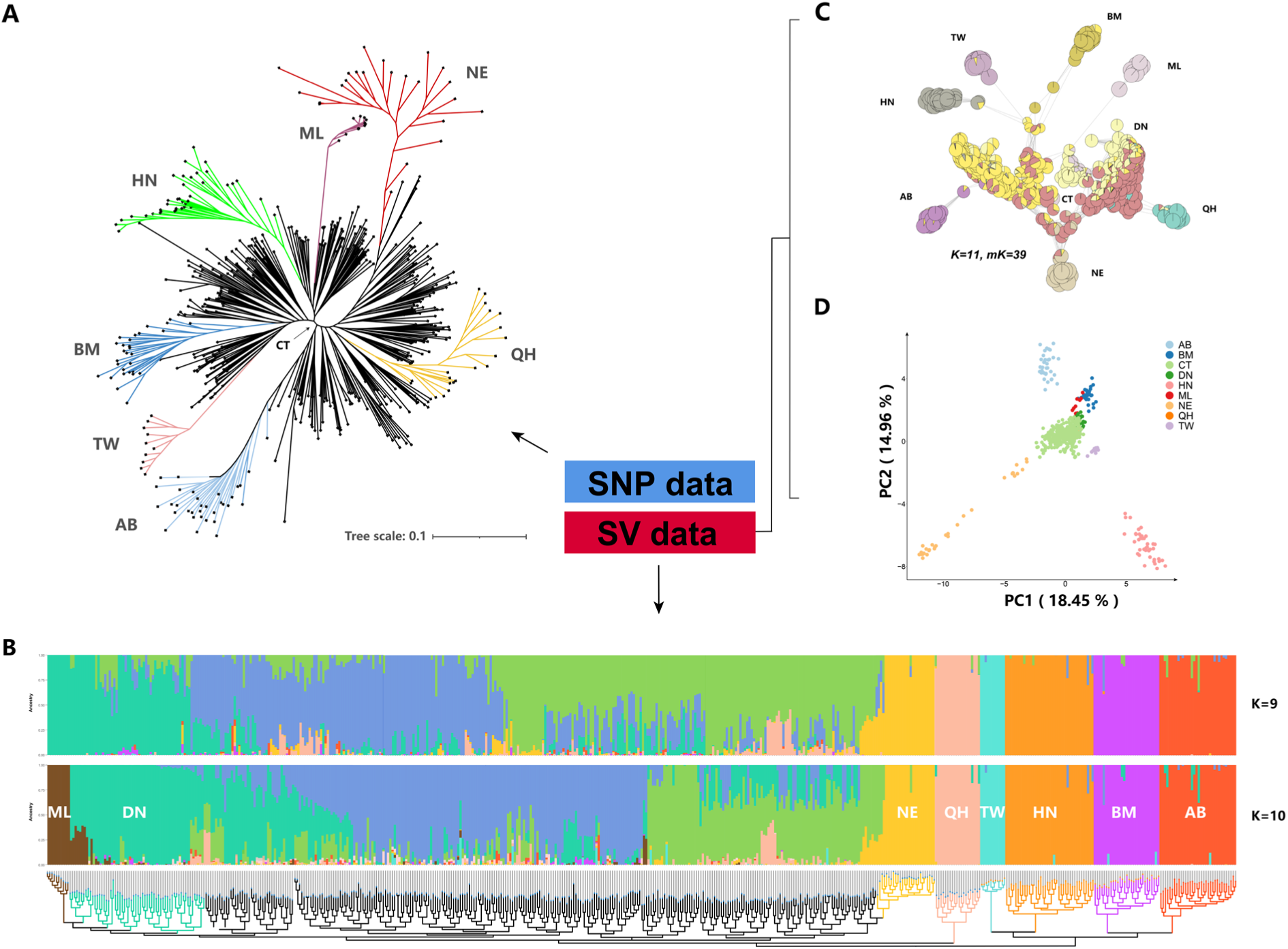
Population structure analysis using SNP and SV data. **A.** Population inferences using SNP data. The eight genetically distinct population groups were divided based on the results obtained from the admixture optimal K value (K=9), which included Aba (AB), Bomi (BM), Central (CT), Hainan (HN), Northeast (NE), Qinghai (QH), Taiwan (TW) and a subpopulation, Malay (ML). **B**. Population inferences using SV data (removing SVs labelled single). The top half of the figure shows the ancestry scores inferred by ADMIXTURE software at K = 9 and K = 10. When K=10, a new *A. cerana* group south Yunnan (DN) was present. The bottom of the figure is the maximum likelihood phylogenetic tree estimated by IQ-TREE using the same data. **C**. All individuals were clustered by state isolation (IBS) matrix using the visualization pipeline Netviewr. Presented here are the results for mk=39, combined with the K=10 file in ADMIXTURE software to plot a pie chart of mixing proportions for each individual sample. **D**. Principal component analysis (PCA) plot using SV data, with the first two components as the X-axis and Y-axis, respectively.

### SVs are genetic variations independent of SNPs, and some are potentially functional

To better understand the contribution of SVs to *A. cerana* genetic variation, we further explored the relationship between SVs and SNPs. Surprisingly, only 13.24% of SVs showed high linkage disequilibrium (LD) with nearby SNPs **(Fig. 6 A)**, which was lower than those found in other species [14, 39]. Additionally, we found that the degree of LD with SNPs was highly dependent on the category of SVs. For example, more than 20% of insertions were highly linked to nearby SNPs, but only 2.23% of inversions were linked with flanking SNPs **(Supplementary Fig. 11 A-C)**. We found a similar relationship between SVs and Indels, suggesting that SVs are a source of genetic diversity that cannot be fully captured by SNPs and Indels **(Fig. 6 A)**.

**Fig. 6:**
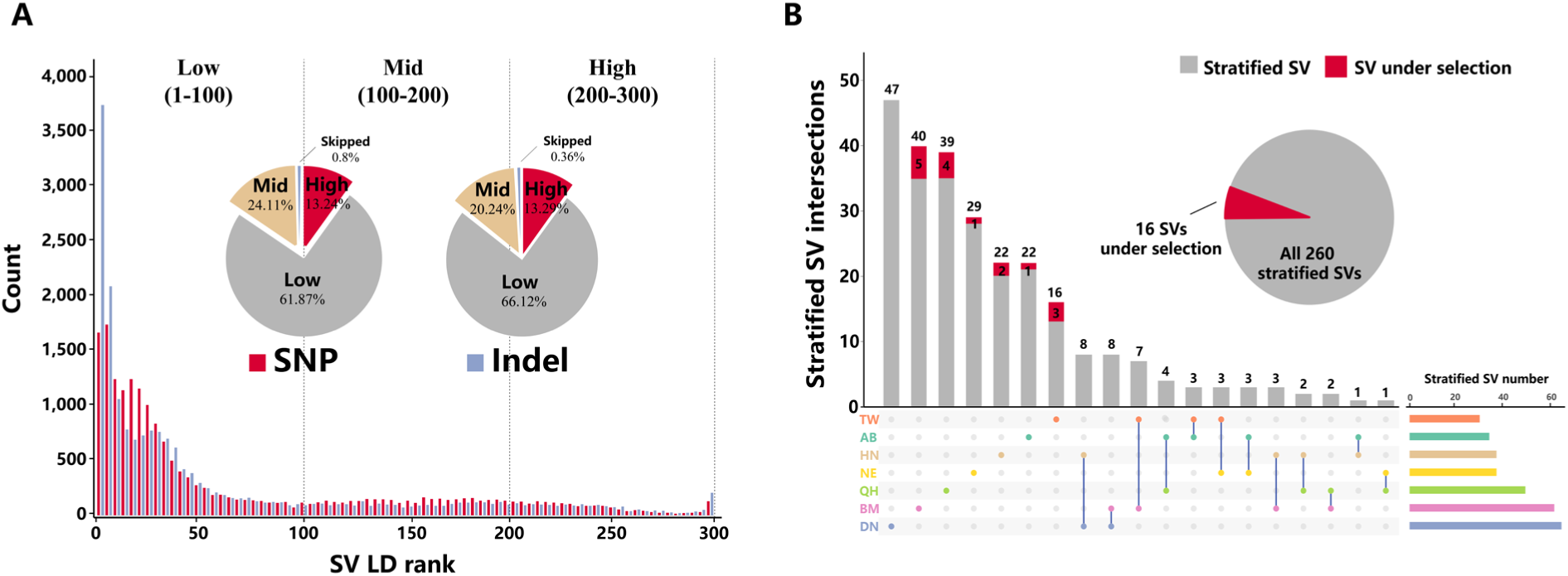
SVs among different *A. cerana* populations. **A.** Histogram of r2 rank number distribution for common SVs (SVs labelled “single” were removed). The graph is divided into SNP-base linkage disequilibrium (LD) rank values and Indel-base LD rank values, which represent the statistics of the 300 SNPs or Indels on either side of the SV whose r2 values exceed the median value, respectively. Note that SVs with fewer than 150 SNPs or indels on either side were not included in this analysis. **B**. The distribution of structural variation in the peripheral population stratified relative to the central population, and the SNP flanking the SV breakpoint was used to infer whether the SV was under selection. The bar graph on the right represents the stratified SVs detected for each group (top 1% of *Fst* values), which were calculated separately for each peripheral population, and the SV unique within or shared between peripheral populations is indicated by the bar graph on the upper side.

Population stratification analysis between each peripheral group and the central group based on the fixation index (*Fst*) identified 260 stratified SVs (top 1% of *Fst* values) **(Supplementary Tab. 11-12; Supplementary Fig. 11 D; Fig. 6 B)**, most of which are located in noncoding (intronic or intergenic) regions. Interestingly, 60% of the stratified SVs residing in genic regions (60%, n=145; intronic or exonic regions) were linked to nearby SNPs (26.9%, n=70; highly and medium linked) **(Supplementary Tab. 13)**. Of all stratified SVs, 16 were subjected to selection **(Fig. 6 B)**. One such SV (one 888 bp deletion) was found in the first intron of the RYamide receptor-like (Lkr) gene in the BM group of *A. cerana*, and this gene has been found to be under selection by SNP data and associated with the foraging labour division of *A. cerana* [3]. As highly stratified SNPs in gene coding regions (high *Fst* values) have been shown to be associated with environmental adaptation in honeybees [3, 20], we hypothesize that environmental adaptation in *A. cerana* could be mediated by SNPs and SVs collectively; highly stratified SVs that are not linked with SNPs (48.4%, n=126) could possibly represent independent genetic variants involved in environmental adaptation.

### SVs are involved in the adaptation of *A. cerana* to climate factors

Correlating environmental variables with genomic variants may provide insights into the complex interplay between genetic and environmental factors, as well as the adaptive mechanisms of organisms [39, 40]. To identify SVs that may be involved in the local adaptation of *A. cerana* to climate factors, we performed genome environment association analyses (GEA) using 19 Bioclim variables from the WorldClim v2 database [40] and two software programs **(see Methods)**. Multiple model cross-validation methods were used to identify the top outliers within the species as local adaptive candidates with high rigor to reduce the probability of false positives [41-43]. A total of 218 cross-validated (LFMM + Bayenv2) SVs were detected as being associated with 19 bioclimate factors **(Supplementary Fig. 12; Supplementary Tab. 9)**, and we named such SVs environmentally associated SVs (eSVs), among which 44 eSVs were located in or near the 2 kb flanking sequences of *A. cerana* protein-coding genes **(Supplementary Fig. 13; Supplementary Tab. 13)**.

GO enrichment analysis was performed on genes containing an eSV in their intragenic or flanking sequences, which could be associated with the climate factors “temperature” or “precipitation”, respectively. The results indicated that genes with eSVs associated with the climate factor temperature were enriched in GO terms including “multicellular organismal homeostasis”, “photoreceptor cell development” and “trachea development”, “locomotor behaviour”, “fibroblast growth factor receptor signalling pathway”, and “cardiocyte differentiation”; genes with eSVs associated with climate factor precipitation were enriched in GO terms such as “open tracheal system development”, “respiratory system development”, and “reproductive structure development” (**Supplementary Tab. 14; Supplementary Fig. 13**); all of these GO terms were considered to be highly correlated with bee local adaptation [42, 43].

Among the genes associated with environmental factors, the sodium/potassium-transporting ATPase subunit alpha gene (*Atpalpha*; LOC107997582) was detected to harbour an eSV, a 330 bp deletion in its intron region **(Fig. 7 A, B),** which is associated with the climate factor mean diurnal range (BIO2), and WGS mapping results indicated that the deletion was only present in the Aba group of *A. cerana* (**Supplementary Fig. 15**). We designed PCR primers anchored on the flanking regions of this deletion, and amplification results revealed that this deletion was only present in the Aba group (**Fig. 7 C; Supplementary Fig. 16**), which demonstrated the reliability of our SV-calling pipeline and indicated that this eSV was the result of an independent evolutionary event.

**Fig. 7:**
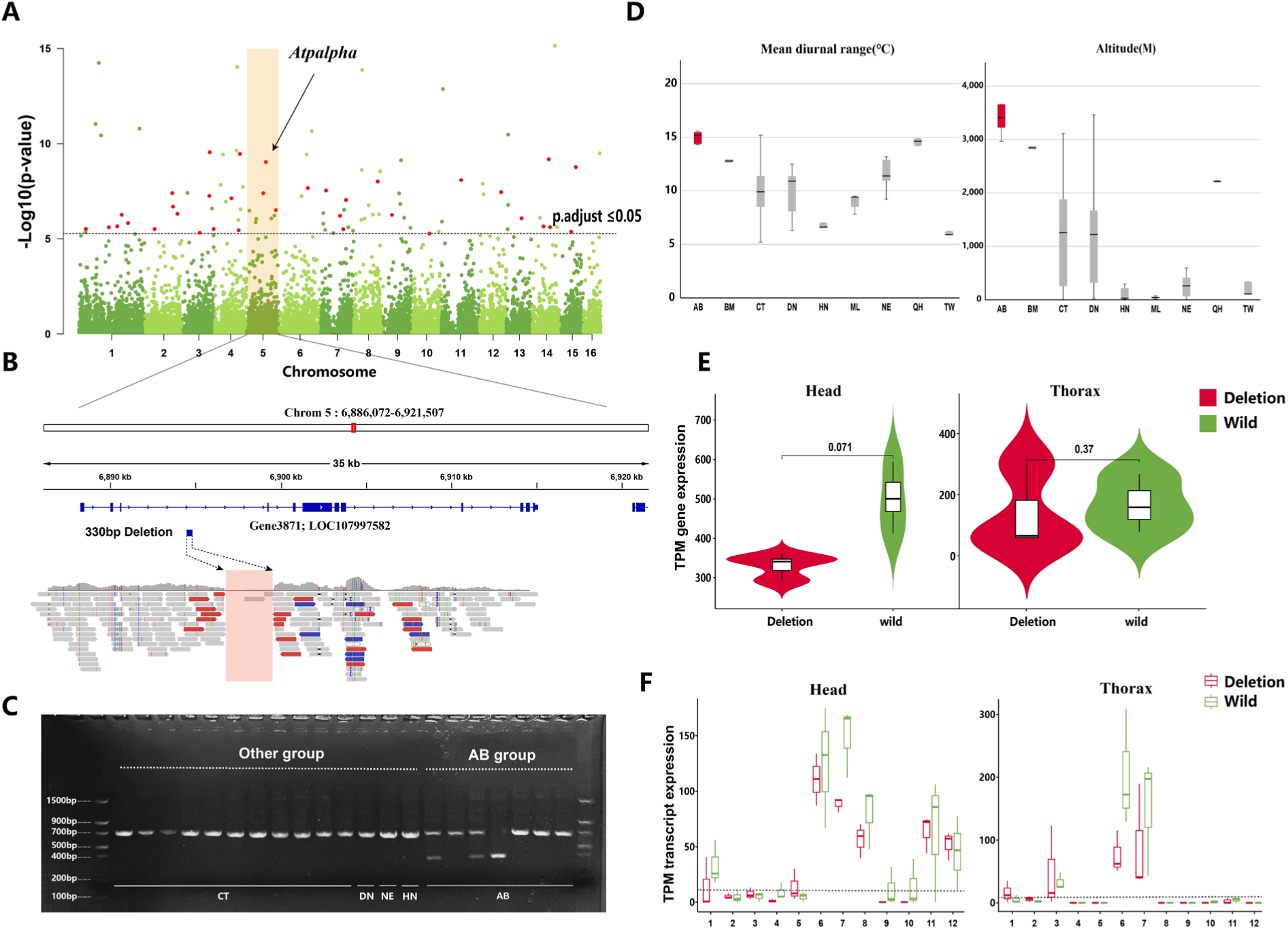
The SV in the *Atpalpha* gene and *A. cerana* cold adaptation. **A.** Manhattan plot of window-based p value statistics. The top panel shows the outlier sites associated with the mean diurnal range (BIO2) bioclimate factors, and the dashed lines indicate the adjusted p value equal to 0.05. eSV sites cross-validated by LFMM and BAYENV2 are marked in red, with eSV in the *Atpalpha* gene being indicated. **B.** A 330 bp unaligned region in the intron of the *Atpalpha* gene was identified in the Aba (AB) group of *A. cerana*. **C.** PCR verification of the 330 bp deletion in the *Atpalpha* gene across different *A. cerana* populations. Longer bands indicate the wild-type *Atpalpha* gene, while shorter bands represent the *Atpalpha* gene with the deletion. **D**. Distribution of mean diurnal range (degrees Celsius) and altitude (meters) across different *A. cerana* populations. **E**. Expression of the *Atpalpha* gene in the brain and thorax tissues of *A. cerana* workers with or without the 330 bp deletion. **F.** Expression of different isoforms of the *Atpalpha* gene with or without the 330 bp deletion in brain and thorax tissues. The dashed line indicates a TPM of 10, and transcripts expressed below this value were considered low expressed.

The *Atpalpha* gene encodes an integral membrane cation anti-transporter, Na+/K+ ATPase (*ATPase*), which plays a crucial role in maintaining ion homeostasis across the plasma membrane using ATP to shuttle Na[+] and K[+] ions [44, 45]. It has been reported that the *Atpalpha* gene exhibits temperature sensitivity and is associated with cold adaptation [46-48]. For example, fruit flies show a decrease in ATPase activity after cold acclimation, and this response is tissue-specific [47, 49]. Similarly, species-specific responses have been observed, indicating certain threshold temperatures beyond which the compensation of the pump is elicited, possibly contributing to the adaptation of these species to different geographical environments [50, 51]. The habitat of the *A. cerana* Aba population has the largest diurnal temperature range, the highest altitude, and the lowest annual mean temperature in summer compared with that of the other populations (**Fig**. **7 C; Supplementary Fig. 17**). Considering the combination of these climatic factors, we hypothesized that 330 bp of SV in the *Atpalpha* gene could be related to the cold tolerance of the *A. cerana* Aba population.

To explore the potential effect of this eSV on the *Atpalpha* gene, we investigated the expression of this gene using transcriptome data obtained from the head and thorax of *A. cerana* workers in the Aba population. We sequenced the transcriptomes of six individuals from the same *A. cerana* colony, including three individuals with the deletion in the intron of the *Atpalpha* gene and three individuals without this deletion. Our findings revealed that the expression level of the *Atpalpha* gene in *A. cerana* workers with this deletion was lower than that in workers without this deletion, particularly in head tissues (**Fig. 7 E, P value=0.071**). In addition, different isoforms of the *Atpalpha* gene were differentially expressed between *A. cerana* individuals with and without this deletion (**Fig. 7 F**). Previous studies have shown that changes in *Atpalpha* gene expression are related to temperature adaptation [46, 51, 52] and that maintaining extracellular ion homeostasis in the brain is vital for cold hardiness in insects [47-49, 53]. Therefore, the eSV, which could alter the expression of the *Atpalpha* gene both qualitatively and quantitively in the brain **(Fig. 7 E**, **Fig. 7 F)**, likely contributes to the cold tolerance of *A. cerana* in the Aba region.

## Discussion

The Asian honeybee, *Apis cerana*, is an important agricultural and economic insect with a widespread natural distribution in Asia [1, 2, 6, 54]. It has been demonstrated that mainland *A. cerana* is composed of one central ancestral group and multiple peripheral groups that radiated from it [3, 17]. Studying locally adapted populations, with higher fitness in local environments, is essential for understanding the evolution and genetic diversity of this species and for predicting the effects of environmental change on the distribution of these honeybees [20, 55, 56]. The unique distribution pattern of *A. cerana* suggests that it could be used as a model to investigate the molecular basis underlying adaptation. In addition, *A. cerana* populations have declined [5-8]; therefore, understanding the adaptive mechanisms of *A. cerana* populations to diverse habitats could provide guidance for the future conservation of this important pollinator.

During the past few years, several large-scale whole-genome resequencing projects have been performed for *A. cerana*, which not only revealed its population structure but also uncovered some genes related to its adaptation [3, 9, 10]. However, only one single reference genome was used in these studies. It was shown that a single reference genome cannot adequately reveal the genetic diversity of one species [11, 13-15, 57]. Therefore, in this study, we utilized PacBio high-fidelity (HiFi) long-read sequencing and high-throughput chromosome conformation capture (Hi-C) techniques to generate a chromosome-scale reference genome sequence for the central ancestral population group of *A. cerana* **(Fig. 1)**. The combination of these advanced sequencing technologies allowed us to overcome the challenges posed by heterozygous sites and obtain a highly accurate and comprehensive reference genome. In addition, we constructed the pan-genome for *A. cerana* based on the reference genome of its ancestral population group and 525 whole-genome resequencing datasets **(Fig. 1)**. Our results revealed that the pan-genome of *A. cerana* is ∼345.2 Mb in length, including 127.5 Mb of sequences that are absent from the reference genome; the pan-genome harbours more genetic information than any single reference genome **(Fig. 1)**. That is, the construction of the pan-genome allows for the considerably more comprehensive genotyping of genetic variations across *A. cerana* populations. Furthermore, we identified 16,587 protein-coding genes in the pan-genome, and 31.32% of them were flexible, i.e. variably present, across *A. cerana* populations; these flexible genes were enriched in biological processes such as response to external stimuli, indicating their potential roles in adaptation **(Fig. 2)**. Mapping of this variable gene content across populations could inform future breeding strategies aiming to increase the fitness of the focal *A. cerana* population.

Previous whole-genome resequencing projects on *A. cerana* mainly focused on small-sized genetic variations, such as SNPs and small Indels; meaning that large-sized variations, such as structural variants (SVs), were largely overlooked [3, 9, 10]. Recently, researchers have revealed that SVs are key contributors to phenotypic variation, which could have important effects on phenotypic traits, disease susceptibility, and adaptive capacity [15, 28, 58, 59]. In this study, we identified and characterized SVs across *A. cerana* populations.

After cross-validation, a total of 19,955 population SV sets were identified **(Fig. 3)**. Most SVs showed low linkage disequilibrium (LD) with nearby SNPs and Indels, suggesting that SVs are a source of genetic variation that cannot be fully captured by small-sized genetic variations, such as SNPs and Indels **(Fig. 6)**. In addition, after performing environmental association analysis between SVs and climate factors, a total of 44 SVs were identified as likely involved in the climate adaptation of *A. cerana*. Moreover, further analysis of one such SV, a 330 bp deletion in the first intron of the *Atpalpha* gene, indicated that this variant likely promotes the cold adaptation of *A. cerana* in the Aba region (Sichuan Province, China) by altering the expression of the *Atpalpha* gene **(Fig. 7)**. Our results map a catalogue of large-sized genetic variants across *A. cerana* populations, associate several of these SVs with climate adaptation, and suggest a mechanism of action for an SV in the *Atpalpha* gene linked to cold adaptation in one *A. cerana* population.

## Conclusions

Through the integrative analysis of multiple genomic datasets including reference genomes and the first *A. cerana* pan-genome, our study provides a comprehensive catalogue of the repertoire of genetic variants of this widespread Asian honeybee. The results highlight the importance of using more reference genomes and genotyping larger-sized genetic variations to extend investigations beyond SNPs and small Indels and reveal additional contributors to the molecular basis of adaptation. These genomic resources and catalogued variants will be valuable for researchers interested in further developing our understanding of the molecular basis of adaptation in *A. cerana* and will lay the foundation for future conservation and management of this important pollinator.

## Materials and Methods

For detailed methods, please refer to Supplementary Materials and Methods. In brief, genomic DNA was extracted from *A. cerana* workers at four sites to create the SMRTbell library, which was subsequently sequenced on the PacBio Sequel platform (Pacific Biosciences). To generate long contiguous sequences (contigs), we employed two different assembly strategies: the CCS schema assembly process and the CLR schema assembly process, tailored to the respective PacBio data schemas. Hi-C sequencing libraries were generated following previously established protocols [60]. We utilized the 3D-DNA pipeline [61] to assemble the long contigs sequences to chromosomal level. The integrity of the genome assembly was evaluated using BUSCO v.4.1.4 [18]. Transposable elements (TEs) were discovered using Extensive De-Novo TE Annotator (EDTA) v.1.9.4 [62]. Protein-coding genes were annotated using the MAKER2 (v2.31.9) pipeline [63], incorporating ab initio gene predictions, transcript evidence, and homologous protein evidence. Gene function was determined through InterProScan 5.53-87.0 [64] and BLASTP software. The genotypes of SNPs and Indels located on the reference genome were retrieved from Genome Analysis Toolkit (GATK, version 4.2.1.0) pipeline [65], and subsequent quality control was conducted using Vcftools v0.1.16 [66]. The pan-genome of *A. cerana* was constructed using a reference-guided assembly approach, as previously reported [14], and gene annotation was performed using MAKER2. Gene presence-absence variation (PAV) information was detected from the mapped bam file using SGSGeneLoss v0.1 software [67] Tajima’s D values for core and flexible genes were calculated using Vcftools, which was also utilized for population genetic diversity analysis. Gene expression was quantified using featureCounts [68] and DESeq2 [69]. Phylogenetic analysis was conducted using IQ-TREE [70], ADMIXTURE v1.3.1 [37], and NetView 1.1 [71]. Genome-wide selective sweep analysis was performed using Vcftools and SweeD v4.0.0 [72] to identify variant sites with selection signals. Detection of structural variants (SVs) in whole-genome sequencing (WGS) data was accomplished using Delly v0.8.7 [73], smoove v0.2.8, and Manta v1.6.0 [74], respectively. Cross-validation was carried out using SURVIVOR v1.0.7 [75]. The formation mechanism of SV was inferred using a previously established algorithm [23, 24, 29, 30], and BEDTools v2.30.0 [76] was employed to construct the background model for assessing SV bias. To infer the potential environmental adaptability of *A. cerana* under different climatic conditions, WorldClim Bioclim (v2.0) [40] data were downloaded as predictors for niche modeling. A multi-model cross-validation approach was employed to identify significant environmental associations. Polymerase chain reaction (PCR) was used to verify the structural variation in the Aba population, and RNA sequencing was performed on the heads and chests of three mutant and wild-type workers. Transcripts were predicted using StringTie v2.2.1 [77], and the expression of the *Atpalpha* gene and each transcript was detected.

## Acknowledgements

This work was supported by the National Natural Science Foundation of China (Grant No. 31971397 and 32270445), the Central Public-interest Scientific Institution Basal Research Fund (Grant No. Y2019XK13 and Y2021XK16) and Swiss National Science Foundation (SNSF) (Grant No. PP00P3_202669 to RMW).

## Author Contributions

C.S., S.D.L., H.H.W., and Y.C.L. conceived and designed the study; Y.C.L., C.S., and H.L.S. were involved in methodology; H.H.W., J.Y., L.S., X.M.Z., D.L.L., and G.Z.W. collected bee samples; Y.C.L. and Q.G.W. collaborated on script compilation for data analysis; Y.C.L. contributed to data visualization; Y.C.L. and C.S. drafted the initial manuscript; Y.C.L. Z.Z.X. and H.H.S. were involved in PCR experiments; S.D.L and F.R.W. was involved in the feeding of bees; K.W. and R.M.W gave guidance in the writing; All authors participated in reviewing and editing the manuscript.

## Data Availability

The data generated in this study, including raw sequence data and genome assemblies, have been submitted to GenBank at the National Center for Biotechnology Information (NCBI), under BioProject number <u>PRJNA869845</u> and <u>PRJNA806528</u>, respectively. Custom scripts for conducting the analyses are available at GitHub at the following link: https://github.com/Liyancan233/A.cerana-Pan-genome.

