## Supplementary Fig for "Pan-genome analysis highlights the role of structural variation in the evolution and environmental adaptation of *Asian honeybees*"

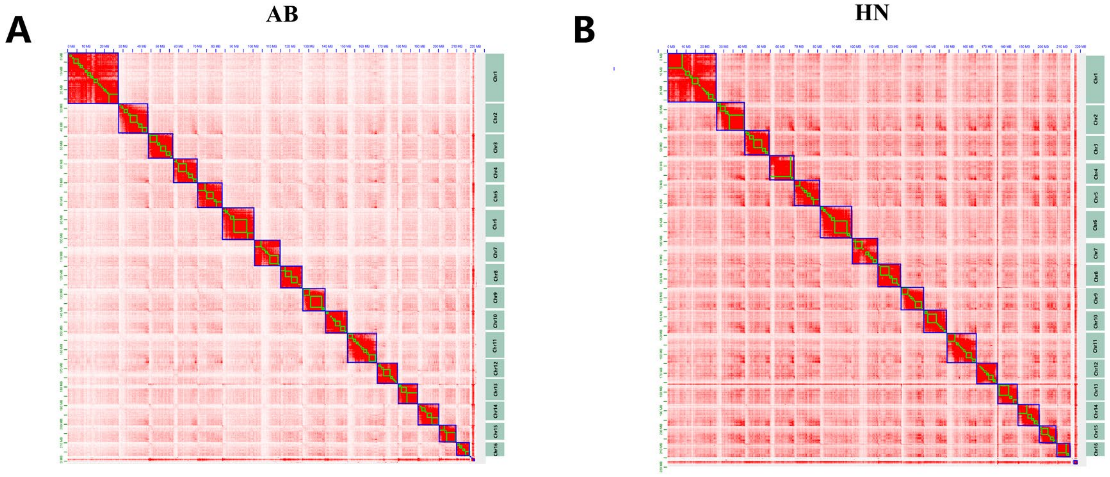


**Supplementary Fig. 1 | Chromosome-level assembly of three *Apis cerana* species.**

Preliminary assembled contigs of three *Apis cerana* were loaded onto chromosomes with Hi-C data. Heatmaps are visualized by Juice box at 2.1-Mb resolution to display the whole genome contact matrices. The order and orientation of the 16 chromosome-scale scaffolds were adjusted using the Amel_HAv3.1 genome as the reference. **(a)** and **(b)** are HB, AB, and HN, respectively.

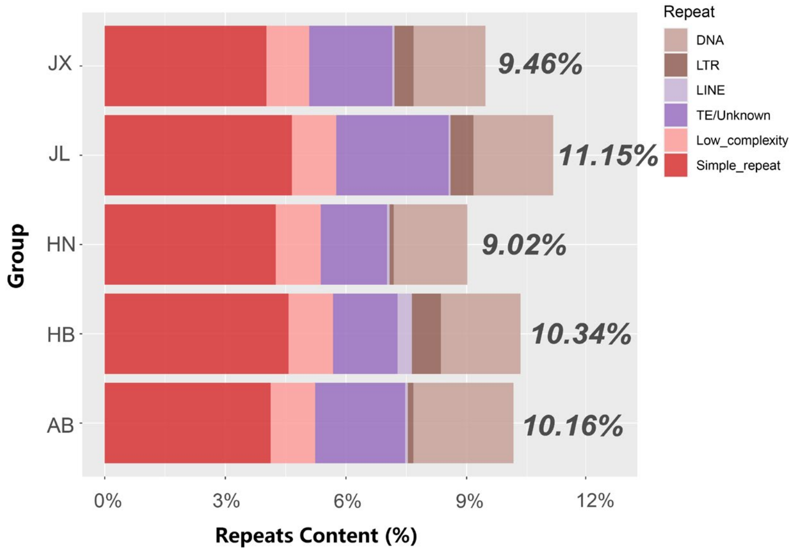


**Supplementary Fig. 2 | The cumulative percentage of repetitive elements for all assemblies.** The x-axis represents the corresponding percentage of genomic repeats. Each type of repetition is indicated by its corresponding colour.

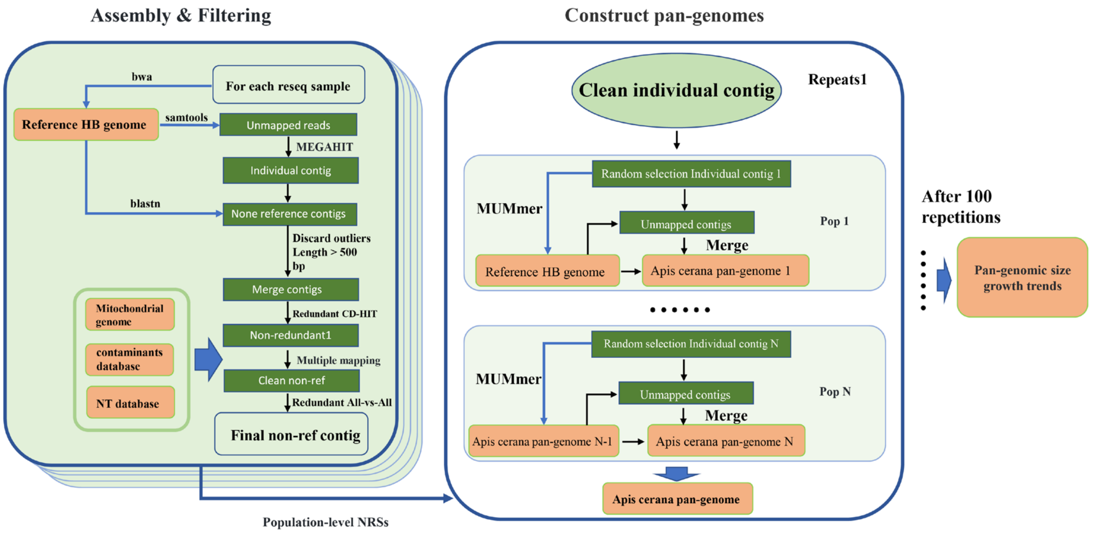


**Supplementary Fig. 3 | Pipeline for pan-genome construction and filtering steps of the *Apis cerana* pan-genome based on unmapped reads with MEGAHIT assembly.**

The WGS reads of each *Apis cerana* accession were trimmed, de-duplicated and then aligned to the reference genome to obtain the nonreference reads (NRRs). The NRRs were assembled and then mapped again to the reference genome to obtain the nonreference sequence (NRSs). The nonreference sequence was then filtered out of contaminants, and redundant sequences were also removed. Clean individual nonredundant sequences are iteratively added into the final pan-genome sequence in a random order method. After 100 replicates, the pan-genome growth trend was obtained, and the assembly with the largest size/least number of contigs was selected as the selection criterion for the final pan-genome sequence.

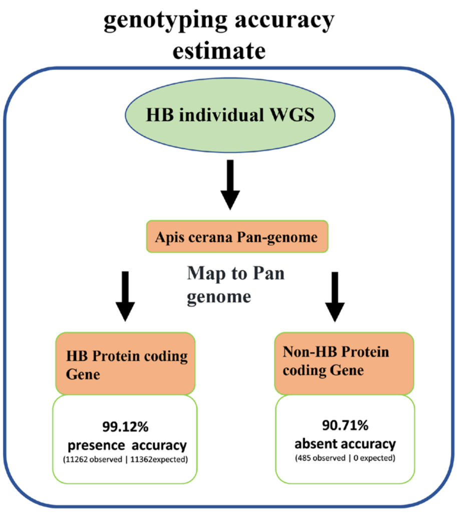


**Supplementary Fig. 4 | Validation of the gene PAV.**

The accuracy of gene presence and absence of variation (PAV) genotyping was estimated by PAV genotyping results from the same region as the reference gene (HB), which was expected to have 100% reference gene presence and 100% nonreference gene absence.

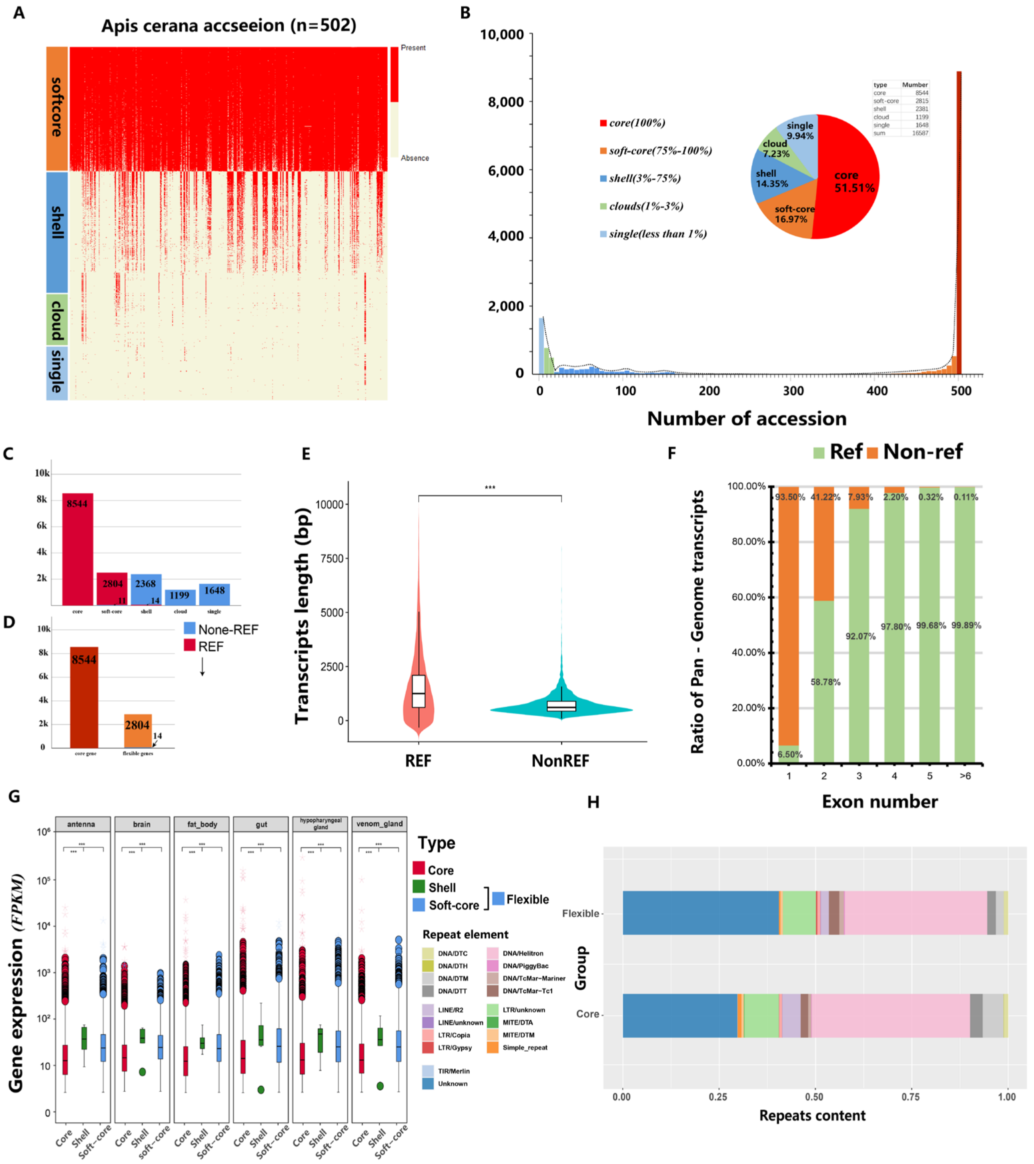

**Supplementary Fig. 5 | Comparison of reference genes and nonreference genes.**

**(a)**. Heatmap of 502 *Apis cerana* accessions showing the presence and absence of variable PAVs. **(b).** Gene number and presence frequency in *A. cerana* pan genes. The pie chart corresponds to the core (present in all accessions), softcore, shell, cloud and single genes. Accessions with low depth (< 5) were excluded from further PAV analysis. **(c).** Distribution of reference genes and nonreference genes **(d).** Core gene and flexible gene in the reference genome. **(e)**. Comparison of exon number between reference and nonreference. **(f)**. Transcript length of reference and nonreference. The Wilcoxon rank-sum test was used for significance analysis (P < 1×10-3). Transcripts longer than 10 kb are not shown in this violin. **(g)**. Expression levels of core, softcore and shell genes in multiple tissues of the *Apis cerana* chromosome genome. From left to right are the antenna, brain, fat body, gut, hypopharyngeal gland, and venom gland. **(h)**. Ratio of transposon elements in the upstream 2 kb of core and variable genes in the *Apis cerana* chromosome genome. The x-axis represents the corresponding repetitive sequence percentage of the genome. Each type of repeat is represented by its corresponding colour.

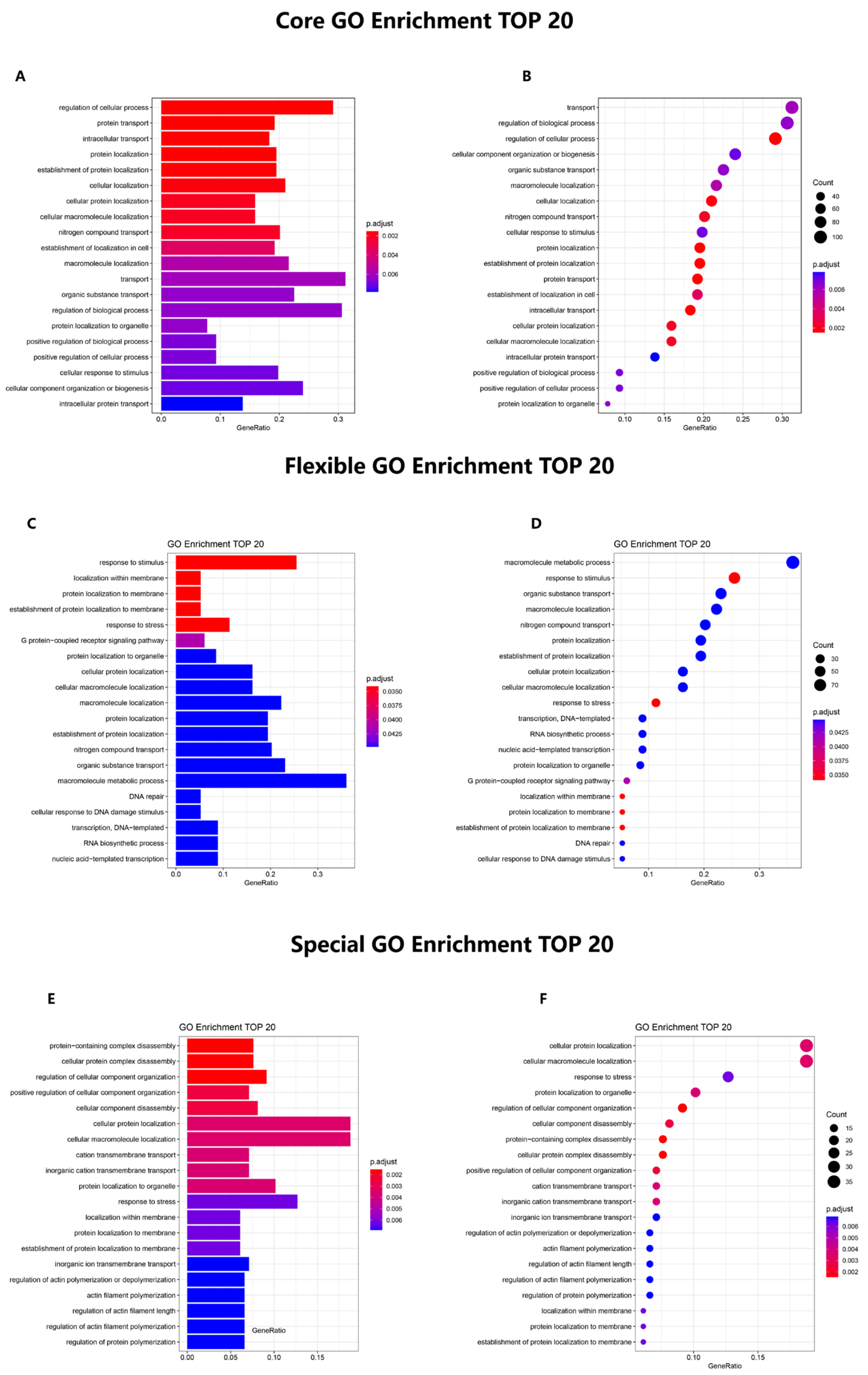


**Supplementary Fig. 6 | GO enrichment of core, flexible and special genes in the *Apis cerana* pan-genome.**

GO enrichment of three classes of genes in *Apis cerana*. Percentage of biological process GO terms for core genes and variable genes. The total number of genes in each GO term and the significance level were calculated by Fisher's exact test. The 20 top GO terms are shown. To better display the data, we drew bar charts and bubble charts for each class, with the top to bottom panels representing gene enrichment of A-B, core gene C-D, flexible gene and E-F, special gene, respectively.

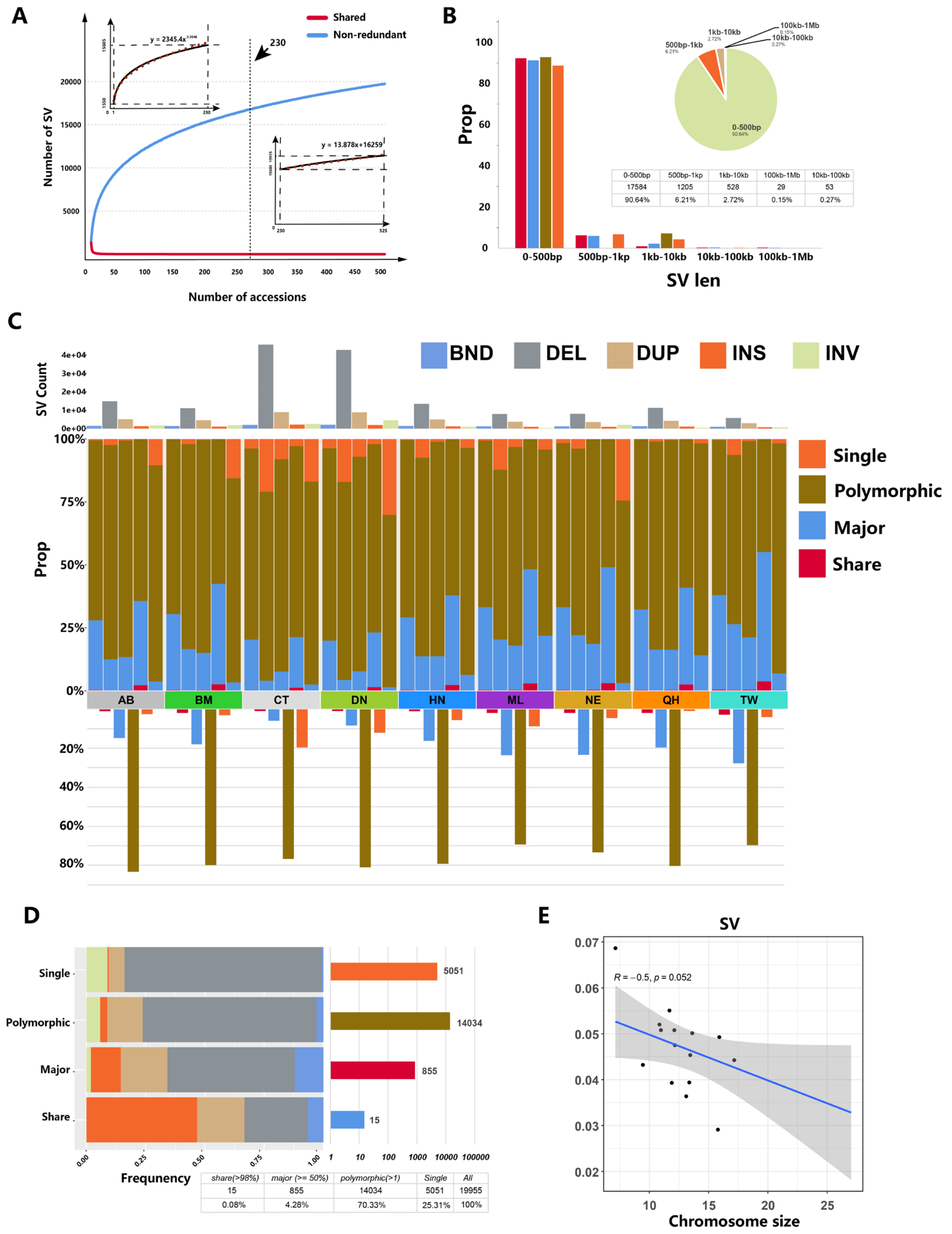


**Supplementary Fig. 7 | Discovery of structural variations in 525 *Apis cerana* accessions.**

**(a).** Discovery of structural variation in 525 *Apis cerana* samples. With the addition of each sample, the newly discovered structural variants (SVs) were gradually merged into a nonredundant set. Shared SVs are shown in red, and nonredundant SVs are shown in blue. **(b).** The distribution of SV length for each shared category, including the shared (identified in all samples), major (identified in ≥ 50% of samples), polymorphic (identified in > 1 sample), and singleton (identified in only one sample) structural variations (SVs), and BND was excluded from this investigation. **(c).** The distribution of SV in different groups, with categorical entries including translocation (BND), inversion (INV), duplication (DUP), insertion (INS) and deletion (DEL), The shared ranks included major SVs (identified in ≥50% of samples), polymorphic SVs (identified in > 1 samples), and singleton SVs (identified in only one sample). **(d).** The number of SVs contained in each category and the proportion of different SV types: translocation (BND), deletion (DEL), duplication (DUP), insertion (INS) and inversion (INV). **(e).** Correlation of chromosome length with nucleotide diversity of SV (from genome-wide resequence data of 525 accessions).

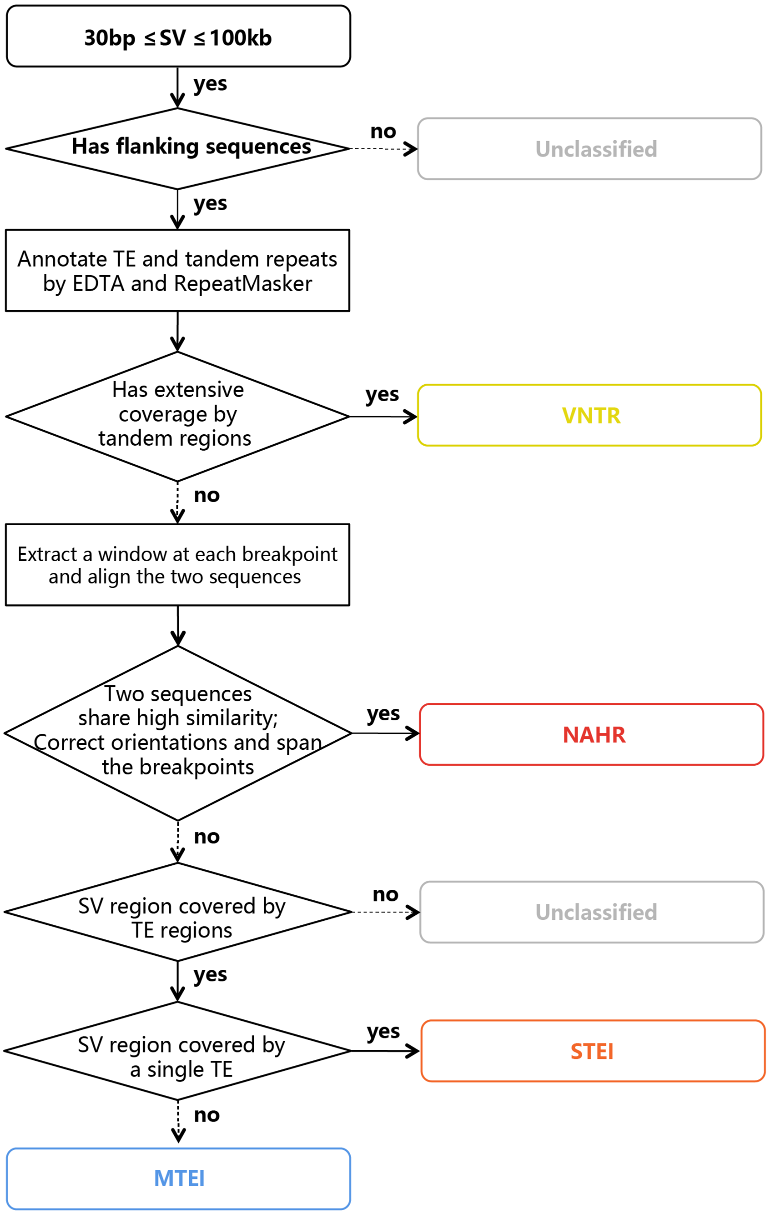


**Supplementary Fig. 8 | Pipeline for classifying SV-formation mechanisms.**
We employed a simplified algorithm for SV classification. Initially, SVs exhibiting over 50% overlap with tandem repeats, simple repeats, or low complexity repeats were categorized as expansions or contractions of VNTRs. Next, SVs were classified as NAHR if the 200bp sequence flanking the breakpoint displayed over 80% identity, indicating homology in the flanking sequence. Lastly, SV sites that overlapped with transposable element (TE) regions were defined as TE-mediated mechanisms for SV. Based on the type of overlapping TE, they were further categorized as single transposable element (STE) or multiple transposable element (MTE).

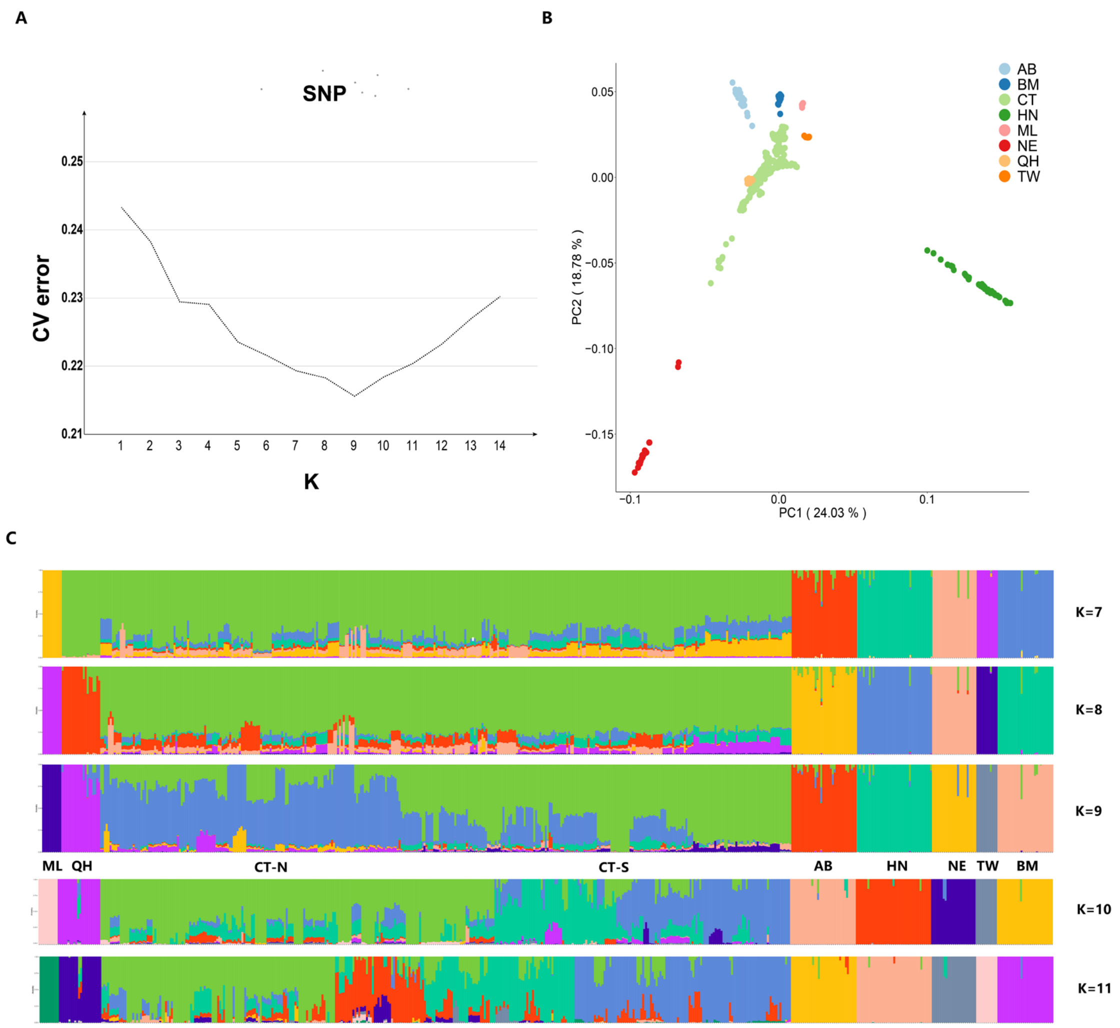


**Supplementary Fig. 9 | Analysis of population structure using SNP data.**

**(a).** ADMIXTURE estimation of the number of groups for K values ranging from 1 to 14, graph showing the best value of *K*=9. **(b)**. Principal component analysis (PCA) plot, with the first two components as the X-axis and Y-axis, respectively. **(c)**. The ancestry scores inferred by ADMIXTURE software at K = 7 to K = 11. Each individual is represented by a single column divided into K colored segments, where K is the number of clusters assumed with lengths proportional to each of the K inferred cluster.

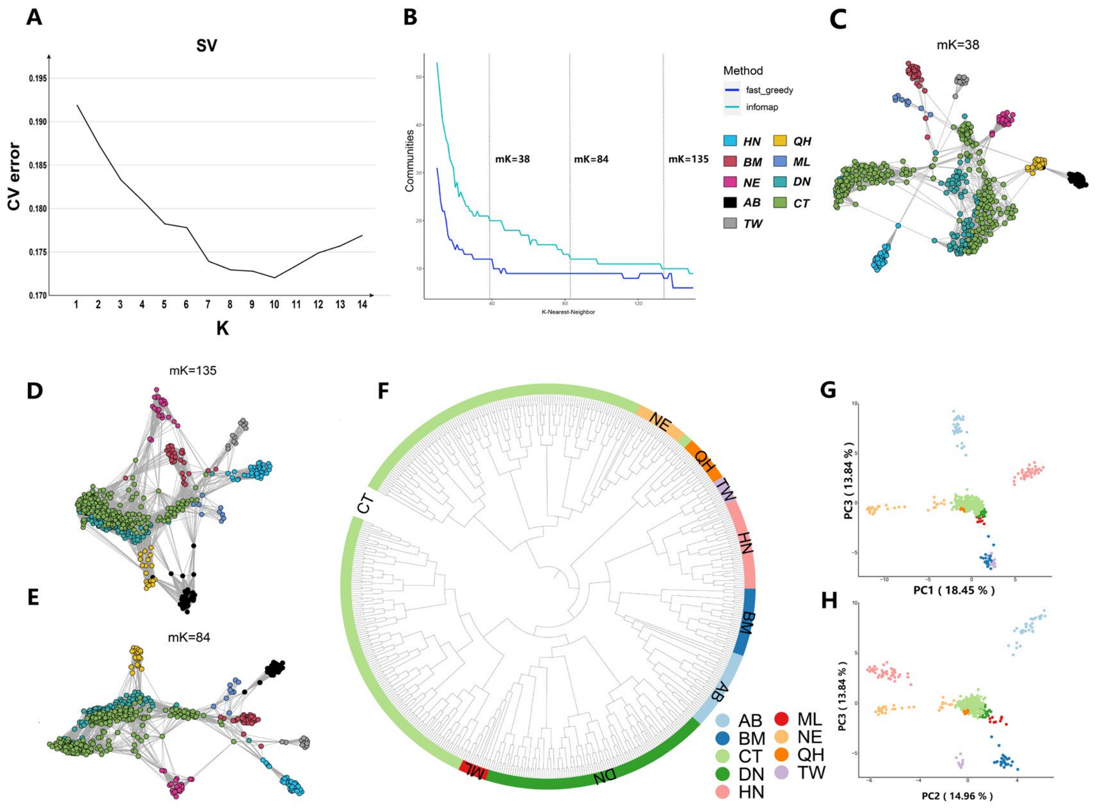


**Supplementary Fig. 10 | Analysis of population structure using SV data.**

**(a)**. Cross-validation of K = 1 through K = 14 clusters from ADMIXTURE analysis with SV data, graph showing the best value of K=10. **(b)**. The number of detected clusters (N) with different numbers of mutual nearest neighbors (mk) is shown using both fast greedy and infomap methods. Networks display the minimum-spanning tree (MST) of mk values at each main inflection point. The **(c–e)** network depicts the minimum spanning tree (MST) of mk values at significant inflection points, highlighting the population structure characteristics at mk=38, mk=135, and mk=84, respectively. **(f)**. The phylogenetic tree estimated by maximum likelihood analysis, obtained by IQ-TREE software using the SV data. **(g-h)**. Principal component analysis (PCA) plot, with the first three components as the X-axis and Y-axis, respectively.

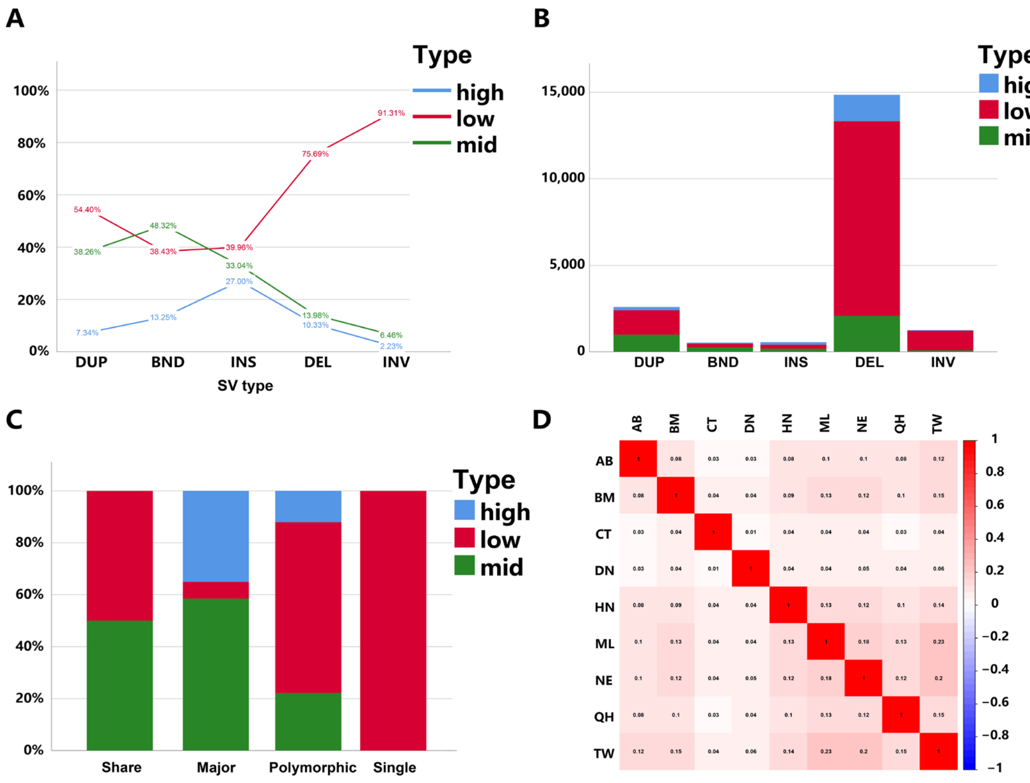


**Supplementary Fig. 11**

**(a)**, **(b)** and **(c)** show the degree of linkage of different classes of SVs to nearby SNPs, respectively.

**(d)** F*st* values between different populations.

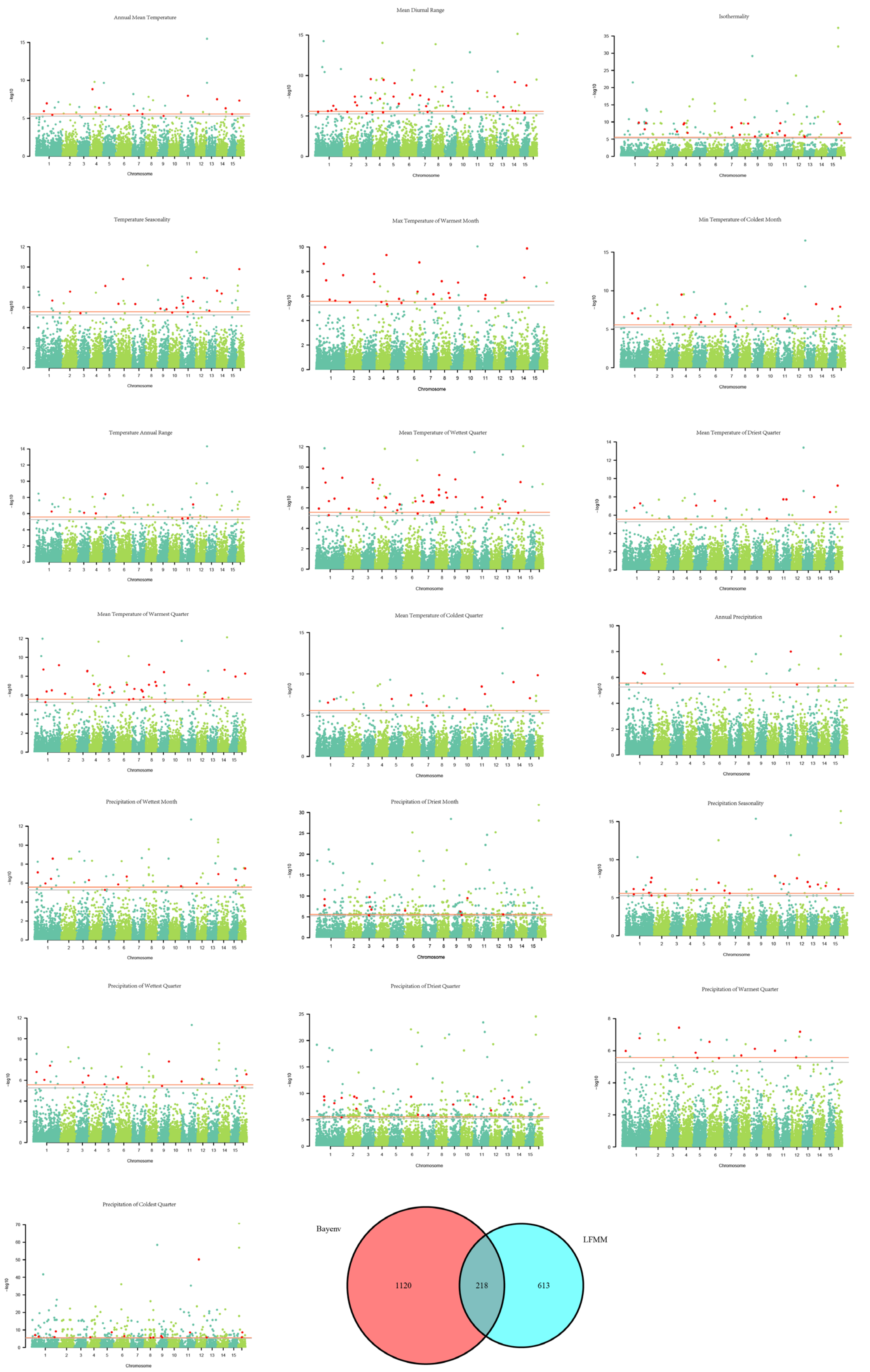


**Supplementary Fig. 12 | Cross-validation method was used to analyze the high environmental**

**correlation SV loci of *A. cerana*.**

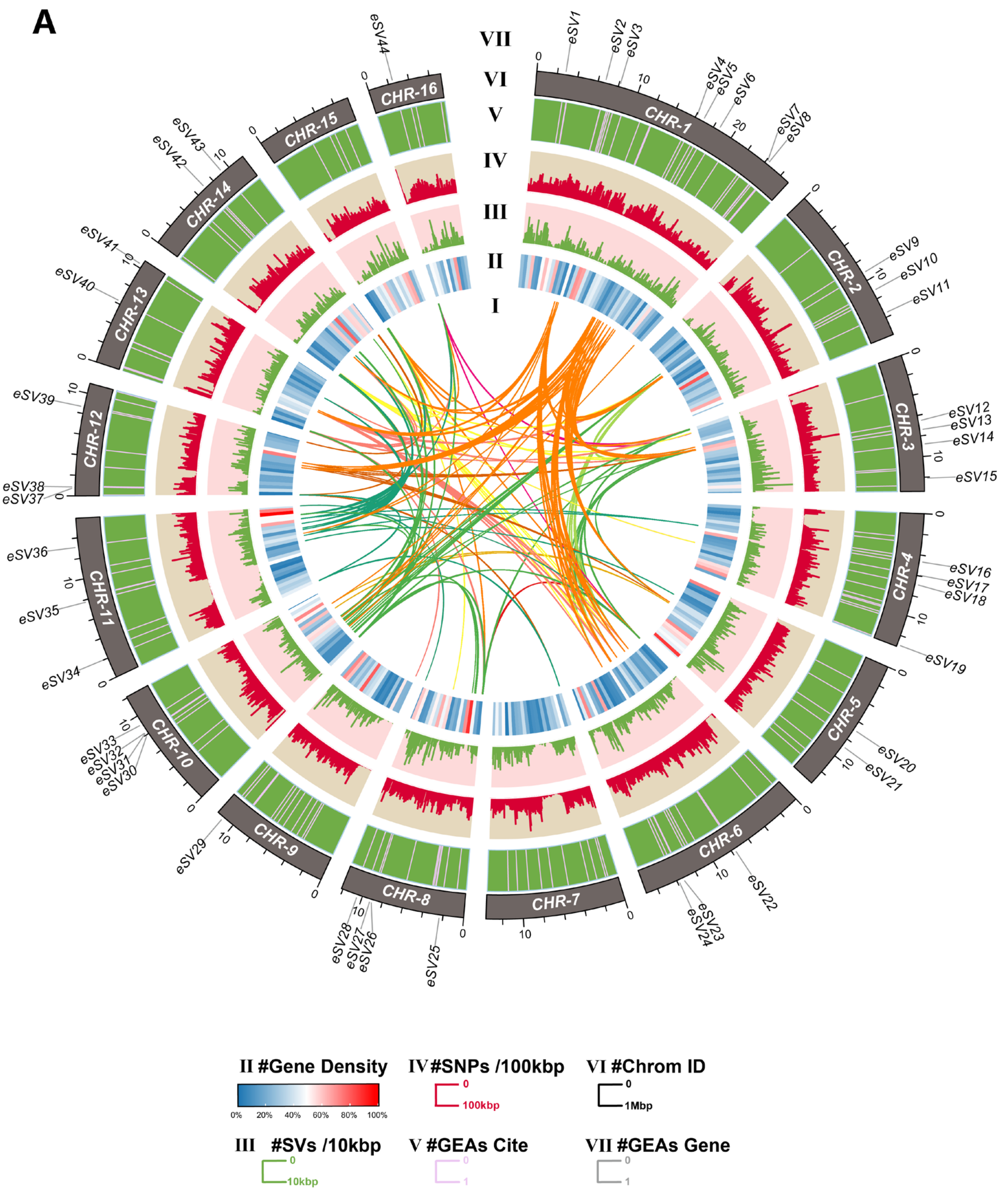


**Supplementary Fig. 13 | The genomic variation landscape of *A.cerana* genome.**

Circos plots show the genomic homology regions, the variation information captured within the population and the highly outlier signal sites of structural variation associated with environmental factors. I-VII, Circos plot showing homologous regions from between pathways to outside pathways (I), gene density (II), SV loci (bin size 100 Kbp) (III), SNP loci (bin size 10 Kbp) (IV), highly outlier eSV loci associated with the environment (V), chromosomal information (VI), and environment-related genes, in which 44 eSV loci occur (before redundancy removal, the number was 81, and one SV may have signal in different Bioclim factors) (VII).

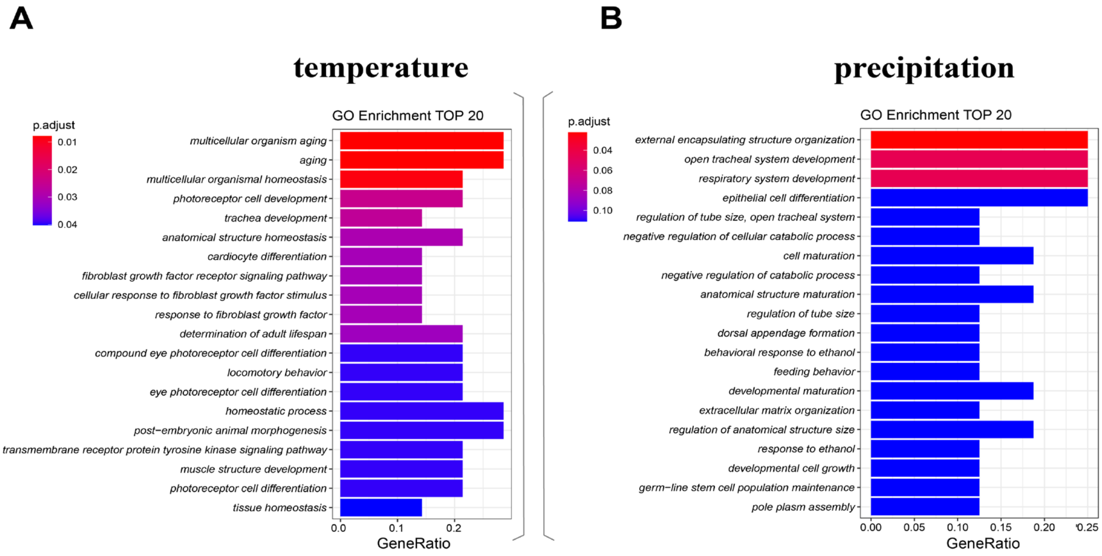


**Supplementary Fig. 14 | GO enrichment analysis of genes in which environmentally relevant SV loci are located.**

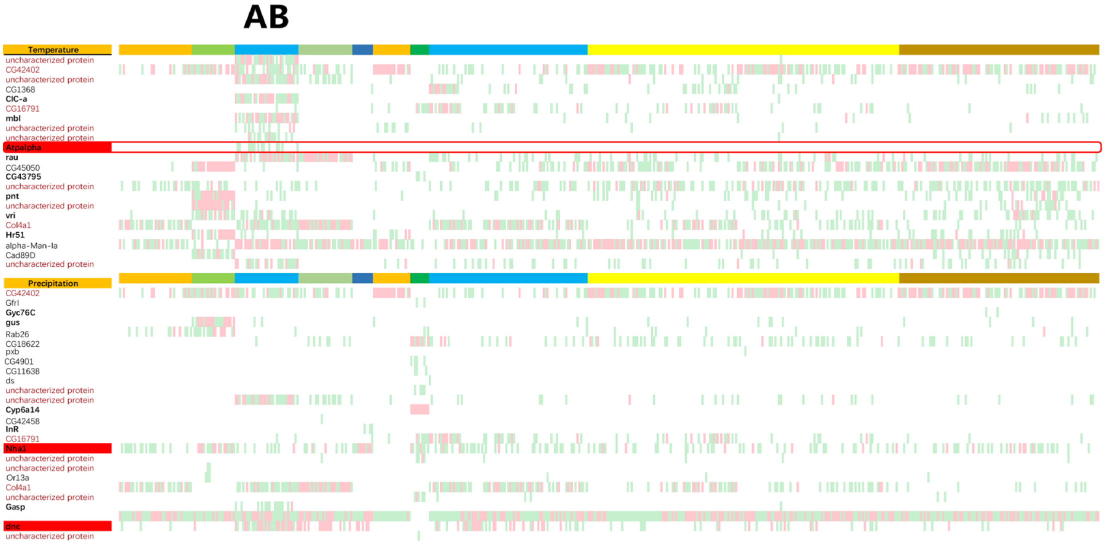


**Supplementary Fig. 15 | SV genotyping data demonstrate the distribution of 44 environmentally associated SVs within different populations.**

The colors in the figure indicate individuals exhibiting mutant characteristics: red represents homozygous mutations, while green represents heterozygous mutations.

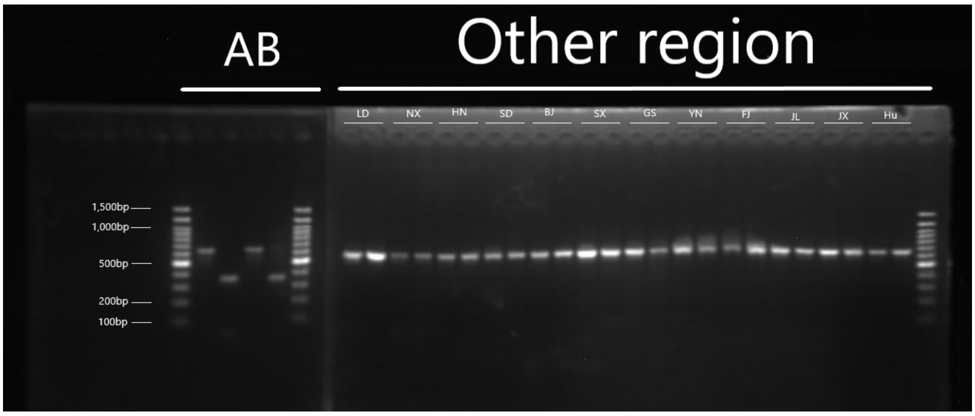


**Supplementary Fig. 16 | PCR verification of the 330 bp deletion in the *Atpalpha* gene across**

different *A. cerana* populations.

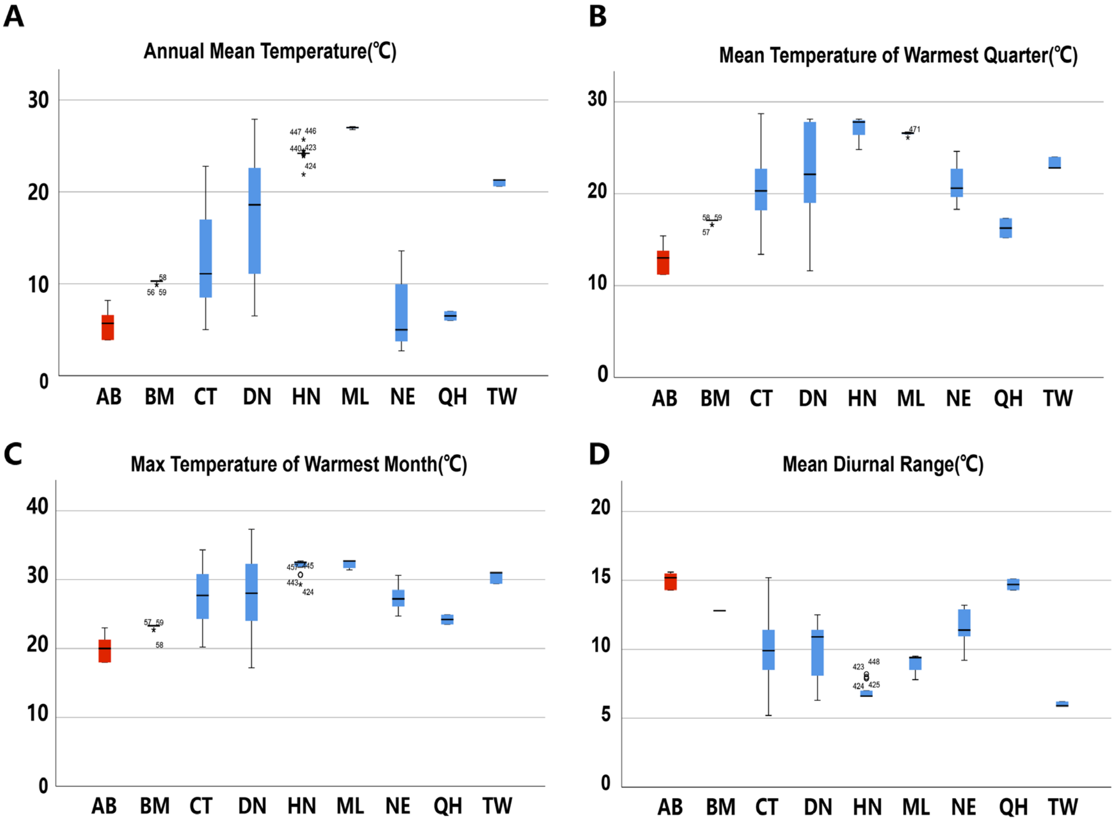


**Supplementary Fig. 17 | Special environmental information for the AB region.**
