## Supplementary Materials and Methods for "Pan-genome analysis highlights the role of structural variation in the evolution and environmental adaptation of *Asian honeybees*"

**Authors:**

Yancan Li[1,3#], Jun Yao[1,3#], Huiling Sang[1,3#], Quangui Wang[1], Long Su[4], Xiaomeng Zhao[5], Zhenyu Xia[1,3], Feiran Wang[1,3], Kai Wang[1,3], Delong Lou[6], Guizhi Wang[7], Robert M. Waterhouse[8], Huihua Wang[1*], Shudong Luo[1,3*], Cheng Sun[2*]

**Affiliations:**

[1]State Key Laboratory of Resource Insects, Institute of Apicultural Research, Chinese Academy of Agricultural Sciences, Beijing, China

[2]College of Life Sciences, Capital Normal University, Beijing, China

[3]Western Research Institute, Chinese Academy of Agricultural Sciences, Changji, China

[4]Institute of Plant Protection, Shandong Academy of Agricultural Sciences, Jinan, China

[5]College of Animal Science, Shanxi Agricultural University, Shanxi, China

[6]Shandong Provincial Animal Husbandry Station, Jinan, China

[7]Department of Animal Science, Shandong Agricultural University, Taian, China

[8]Department of Ecology and Evolution, University of Lausanne, and SIB Swiss Institute of Bioinformatics, 1015 Lausanne, Switzerland

[#]**Authors contributed equally.**

[*]**Corresponding authors:**

### 1. Collection of genomic data in *Apis cerana* individuals

#### 1.1 PacBio, HIC library construction, and sequencing

Prior research have reported that *A. cerana* consists of multiple peripheral groups that are independently derived from a central group (Ji et al., 2020). In this study, we collected worker bee samples from different regions in China, including Shennongjia, Hubei Province (HB), and Yichun, Jiangxi Province (JX), which are considered central regions. Similarly, samples were obtained from Jilin Province (JL) in the Northeast Region and Aba in Sichuan Province. The SMRTbell library was constructed as previous described (Pendleton et al., 2015) and sequenced on the PacBio Sequel platform (Pacific Biosciences). For all PacBio sequencing, the pupae intestines were removed to avoid intestinal microbial contamination. Two modes were employed for the mentioned samples. The Pacific Biosciences Sequel II platform's cyclic consistency sequencing (CCS) mode was utilized for HB and JL to generate HiFi data. Conversely, for JX and AB, CLR sequencing was conducted using the Continuous Long Read mode (CLR) pairs on the same platform. The outcome was 79.8 Gb and 74.3 Gb of HiFi data for HB and JL, respectively, while the CLR mode produced 30 Gb and 31.9 Gb of PacBio long reads for JX and AB, respectively **(Supplementary Tab.2).**

Adequate worker bees (female, diploid, n > 15) were collected in Hubei and Aba regions for HIC sequencing preparation. Following digestion with the MboI enzyme, we utilized the previously described Hi-C library preparation protocol (Belton et al., 2012). The Hi-C library was then sequenced on Illumina NovaSeq 6000 using the PE150 sequencing strategy. To enhance our data, we acquired the published Hainan Region (HN) PacBio CLR and Hi-C data, both of which were generated through drone pupae sequencing and sampled from a single colony. (PRJNA579740) (Wang et al., 2020).

#### 1.2 Short-read data aggregation

The detailed information of the whole genome shotgun reads (WGS) dataset including the sequencing read sizes, and geographic or pedigree information for *A. cerana* pan-genome (APG) was listed before (**Supplementary Tab.1**). In short, most of our data comes from previously published studies, including 180 *A. cerana* from China and 306 from China and neighbouring regions (C. Chen et al., 2018; Ji et al., 2020). In addition, to supplement the data set, we additionally generated 39 resequencing data from 13 sites in 11 administrative provinces in China, including Baoji and Xi 'an in Shaanxi Province, Longyan in Fujian Province, Qingyang and Longnan in Gansu Province, Xinglong in Hebei Province, Haikou in Hainan Province, Changde in Hunan Province, Jilin Jilin in Jilin Province, Yichun in Jiangxi Province, Guyuan in Ningxia, Zibo in Shandong Province and Pu'er in Yunnan Province, with three samples collected at each site. To avoid the effects of potential genetic mixtures caused by the introduction of alien populations, we only collected the worker bees only in local nature reserves. These samples were sequenced to 150 bp paired-end reads with an average coverage of 40x using the Illumina HiSeq X Ten platform. To sum up, Our *A. cerana* data are dominated by China, supplemented by North Korea, South Korea, Thailand, Vietnam, Thailand, Malaysia, Singapore, it covers most of mainland Asian honeybee *A. cerana* habitat.

#### 2. De novo assembly, annotation, and evaluation

##### 2.1 Genome assembly

We acquired PacBio SMRT long reads from five *A. cerana* sites (Aba/Sichuan, Hubei, Hainan, Jilin, and Jiangxi) and employed two de novo assembly approaches. For the HiFi data JL and HB, an initial assembly was created using hifiasm (v0.16) (https://github.com/chhylp123/hifiasm) with the parameter "-l0". Subsequently, Purge_Dups (v.1.01) was utilized to eliminate potential redundant regions in the assembly, using default parameters and only the primary haplotype was retained for each accession.

For long reads generated in PacBio CLR mode, we assembled the three genomes AB, HN, and JX using two different assembly strategies. After self-correction of raw data using Canu v.1.6 (Koren et al., 2017)(https://github.com/marbl/canu) with parameter "-correct", and then performed assembly using Canu and Falcon-1.3.0 (Chin et al., 2016) (https://github.com/PacificBiosciences/pb-assembly), respectively. For Falcon, we set the important parameter "length_cutoff_pr to=8000, ovlp_HPCdaligner_option=-k24-h1024-e.95-l1800-s100, overlap_filtering_setting=--max-diff 100 --max-cov 150 --min-cov 3". Canu v.1.6 used the parameters "minReadLength=2000", which were data-optimized parameters for our data.

After obtaining preliminary assembly, we remapped self-corrected reads to them using minimap2 and performed the three rounds polish on the resulting contigs using Racon (Vaser et al., 2017)(https://github.com/isovic/racon). Next, we applied Pilon v.1.23 (Walker et al., 2014) for three rounds of contig polishing using available resequencing data. Potentially redundant regions in assembly are excluded using default parameters by running Redundans (Pryszcz & Gabaldon, 2016), and the unique sequences in the Falcon assembly that were not included in Canu were integrated into the final genome **(Supplementary Tab.2)**.

##### 2.2 Pseudo-chromosome construction

Chromosome level assembly was performed using the 3D-DNA pipeline (Dudchenko et al., 2017). In brief, for AB, HB, and HN, raw Hi-C reads were trimmed and filtered by fastp (S. F. Chen et al., 2018) with the default parameters. The processed Hi-C reads were aligned to the polished contigs using the Juicer pipeline (Durand et al., 2016). After manual correction using Juicebox Assembly Tools (Durand et al., 2016), The contig were ordered and oriented to 16 pseudomolecules **(Supplementary Fig.1; Supplementary Tab.3)**. The remaining assemblies (JL and JX) were anchored and oriented to chromosomes using the reference-guided software RagTag (v2.1.0) (Alonge et al., 2022) with the default parameters.

##### 2.3 Transposable element (TE) annotation

TE elements were discovered using Extensive De-Novo TE Annotator (EDTA) v.1.9.4 (Ou et al., 2022), which combined the raw predictions of long terminal repeat (LTR), Terminal-inverted repeat (TIR), and Helitron elements produced by LTRharvest_parallel (v1.5.10) (Ellinghaus et al., 2008), LTR_FINDER_parallel (v1.0) (Ou & Jiang, 2019), LTR_retriever (v2.6)(Ou & Jiang, 2017), Generic Repeat Finder (v1.0)(Shi & Liang, 2019), TIR-Learner (v1.23) (Su et al., 2019), and HelitronScanner (v1.0)(Xiong et al., 2014). After extra basic and advanced filters, the remaining TE sequences were identified by RepeatModeler (v2.0.1)(Flynn et al., 2020). RepeatMasker (v4.1.0) (Chen, 2004) finally performed homolog annotation with the TE library (EDTA.TElib.fa). The non-redundant TE library was combined with a library of simple repeats and satellites generated by RepeatMasker (v4.1.0) to represent the final repeat regions **(Supplementary Fig.2; Supplementary Tab.4)**.

#### 2.4 Gene annotations annotation and evaluation

The protein-coding genes were predicted in Hi-C oriented genomes (AB, HB, and HN) with MAKER2 (v2.31.9) pipeline (Holt & Yandell, 2011). To provide transcript evidence, we obtained previously reported RNA-sequencing (RNA-seq) data (including six tissue data: antenna, brain, fat body, gut, hypopharyngeal gland, venom gland) (Xu et al., 2017) and combined it with our data for further analysis. HISAT2 (v.2.2.1) (Kim et al., 2015) was used to perform splice alignment of RNA reads to the ref genomes (HB) with the "--dta-cufflinks" parameter. Potential transcripts were de novo assembled, using StringTie v2.2.1 (Pertea et al., 2015) with default parameters. Candidate coding regions were identified using TransDecoder v5.5.0 (https://github.com/TransDecoder/TransDecoder ). For ab initio gene prediction, we trained SNAP (v2006-07-28) (Korf, 2004) using MAKER2 and performed AUGUSTUS (v3.2.3) (Stanke & Morgenstern, 2005) prediction using the "honeybee1" model. The sequence was masked against the TE library produced by EDTA (v.1.9.4) in the MAKER2 package with "rmlib". Non-redundant *A. cerana* protein sequences downloaded from the UniProt Swiss-Prot database were passed to MAKER2 as homologous protein sequences along with transcripts compiled by StringTie in the previous step. After two rounds of model training, we ran the final MAKER2 and combined it with the SNAP HMM and AUGUSTUS gene model trained previously to synthesize the final gene annotation. The longest transcript of each predictive gene model was considered representative.

Functional information was added to the "gff" files generated in the previous step using a multi-software annotation pipeline. Briefly, the protein sequences of the predicted genes were compared against the InterPro database using InterProScan 5.53-87.0 (Blum et al., 2021) to identify functional protein domains with parameters "-appl Pfam -iprlookup -pa -goterms" . Gene function was determined by comparing protein sequences with the GenBank non-redundant protein database using BLASTP with the options option "-p BLASTP-E 1e-05-b 5 -v 5 -a 4". Finally, we evaluated the completeness of predicted genic using the hymenoptera_odb10 database (for Hymenoptera species) of BUSCO v.4.1.4 (Simao et al., 2015) with default parameters

### 3. *A. cerana* pan-genome construction and gene annotation

#### 3.1 Pan-genome construction based on short reads data

The pan-genome of *A. cerana* was constructed using a reference-guided assembly approach, as previously reported (Li et al., 2021). WGS data for each accession underwent quality control using fastp 0.23.2 (S. F. Chen et al., 2018) with parameters "-q 5 -n 5 -u 30". The resulting cleaned data were aligned to the reference genome using SAMtools (Li et al., 2009) with parameters "-b-f 4, -f 68-f 8 and -f 132-f 8" to extract unaligned reads.

For each accession, we followed this procedure: The unaligned reads, including all unmapped paired reads and unpaired single reads, were assembled with MEGAHIT (v1.2.9) (Li et al., 2015) using default parameter, resulting in the original contig. After filtering out contigs with the length of less than 500bp, the remaining contigs were aligned against the reference genome using the nucmer

program in MUMmer (v4.0.0) package (Delcher et al., 2002). Contigs that were highly similar to the reference sequence were filtered out (whole contigs with identity > 0.9 and match ratio > 50%). Redundant sequences were also filtered using CD-HIT (v4.8.1) (Fu et al., 2012) with the command "-c 0.9 -G 0 -aL 0.90 -AL 500 -aS 0.9". To remove contaminants, we set up a filtration pipeline:(1) Kraken2 (Wood & Salzberg, 2014) was first used to discard unaligned contigs in human, archaeal, bacterial, and viral genomes and collection of known vectors (UniVec_Core) with standard database (201904). (2) The remaining contigs were the blast to NCBI nt database (20200203) to identify other potential contaminants. (3) The mitochondrial genome (GenBank: NC_014295.1) was also identified and then removed. (4) Finally, the remaining contigs were subject to an all-versus-all alignment with BLASTN (with parameters "-e 1e-05") to ensure non-redundancy.
After completing the above steps, we combined the results for all individuals. To ensure that the merged contigs set was absent in the reference genome, we used nucmer (with parameters "-c 90 -l 40") of MUMmer (v4.0.0) to identify contigs with a match ratio over 90%, and then deleted. The *A. cerana* pan-genome sequences were finally generated by combining the HB reference sequences and the non-reference assembled contigs (final non-reference non-redundant sequences data) **(Supplementary Fig.3; Supplementary Tab.6)**.

##### 3.2 Annotation of pan-genome genes

Non-reference assembled contigs were annotated using MAKER2 (v2.31.9) as previously mentioned (2.4 Materials and Methods). SNAP (v2006-07 07-28) and Augustus (v3.2.3) were used for gene prediction using the trained model. Transcript evidence was obtained from the available ACSN-2.0 transcript data (https://ftp.ncbi.nlm.nih.gov/genomes/all/GCF/001/442/555/GCF_001442555.1_ACSNU-2.0/) and the data predicted in the previous step. ACSN-2.0 protein information and newly predicted gene information were used as protein evidence. Repeat sequences were masked using the TE library generated with EDTA. The result transcripts were aligned to reference transcripts to remove potential redundant redundancies. Predicted proteins with less than 30 amino acids were filtered out. Functional annotation of genes was performed using the methods described above. Specifically, protein functional domains are pre-determined by InterProScan and BLASTP was used to identify gene function. KOBAS (Bu et al., 2021) was used to assign gene ontology (GO) terms and Kyoto Encyclopedia of Genes and Genomes (KEGG).

#### 4. *A. cerana* pan-genome analyses

##### 4.1 Calculation of sequence pan-genome size

The nonredundant non-reference sequence (produced in procedure 3.1) of each accession was used for the calculation sequence pan-genome size. we established a pipeline to study the growth of the *A. cerana* pan-genome size with respect to population size. In brief, we iteratively added each accession one by one to complete the pan-genome in an unordered fashion, and the nucmer program in the MUMmer (v4.0.0) package (Delcher et al., 2002) was used to filter the shorter contigs in the paired comparison with a high degree of similarity (identity over 0.9 and the match ratio more than 80% of the whole contig). After 100 repetitions, we obtained a growth curve of sequence pan-genome size. Pan-genome integrity assessment using samtools with "flagstat" parameter. Five long read reads were mapped to the five high-quality genomes and APG respectively to detect the mapping ratio for the assessment of pan-genome integrity **(Supplementary Tab.7)**.

##### 4.2 Gene presence-absence variation (PAV) and Calculation of pan gene size

To call the gene presence and absence matrix, we mapped the raw datas for each accession to the pan-genome using BWA-MEM v0.7.12 (Li & Durbin, 2009) with default parameters. The gene PAV information was detected from the mapped bam file using SGSGeneLoss v0.1 software (Golicz et al., 2015) with the option "minCov=2 lostcut=0.2". If more than 80% of exon regions were covered by at least 2 reads, this gene was called present with the "1/1" genotype. As lower depth regions may produce potential bias in the acquisition of read evidence (Li et al., 2021) (Golicz et al., 2016), we only retained the sample with average coverage over 0.8 and average depth over 5. After obtaining the gene PAV matrix, the completely random algorithm of PanGP (v.1.0.1) (Zhao et al., 2014) was used to simulate the gene pan-genome size, with the sample size set as 500 and sample repeat set as 30.

##### 4.3 Gene PAV analysis

We defined variable genes and divided them into 5 categories: core, softcore, shell, cloud, and single genes. The softcore, shell, cloud, and single genes were present in 75–100%, 3–75%, 1–3%, and less than 1% of the population, respectively. For the convenience of analysis, softcore and shell genes were combined as flexible genes, while cloud and single were classified as specified population genes. To explore the differences in gene expression levels between different classifications, we aligned RNAseq from different tissues (see 2.4 methods) to the reference genome (HB) using HISAT2. Before aligning, the index of the longest transcript isoform of gene was built based on our annotation, and only one CDs region gene was excluded. The program featureCounts in package Subread 1.6.4 (Liao et al., 2014) was used to summarize read counts on exons with the "-p" parameter. Relative Log Expression (RLE) values were calculated using the R package DESeq2 (Love et al., 2014). Vcftools v0.1.16 (Danecek et al., 2011) was used to calculate Tajima's D by comparing variations of core and flexible genes with default parameters. Genotyping evaluation of the gene PAV matrix was performed using a dataset from the same region as the reference genome, with five randomly selected individuals mapped to the pan-genome. Theoretically, genes located in the reference genome should all be captured and non-reference genes should not be captured **(Supplementary Fig.4)**.

#### 5. Genotyping and Characterizing the *A. cerana* genetic variation map

##### 5.1 Identification of SNPs and indels from 525 accessions

The genotypes of SNPs and indels located on the reference genome (HB) were retrieved from Genome Analysis Toolkit (GATK, version 4.2.1.0) pipeline (McKenna et al., 2010). In detail, raw WGS reads were first filtered using Trimmomatic v0.39 (Bolger et al., 2014) with a default parameter. The clean reads were aligned to the reference genome using BWA-MEM. picard v2.25.7-0 (https://broadinstitute.github.io/picard/;) was further used to filter out duplicate mapped reads and uniquely mapped reads were retained. SNPs and indels were identified using GATK with HaplotypeCaller parameters. After combining the "gvcf" of the resulting single individuals, genotyping was performed using the GenotypeGVCFs function. Here, we only considered variants located in chromosomes, and those located in scaffolds were excluded from further analysis. To ensure high-confidence variants, we conducted two rounds of filtering. the detailed filtering processes were as follows: (1) the variants were first filtered using GATK with "QD < 2.0, FS > 60.0, MQ < 40.0" parameters. (2) then, Secondary filtering was performed using vcftools v0.1.16 (Danecek et al., 2011) with "maf 0.05, max-missing 0.5-min-meandP 2, Max-meandP 40, hwe 0.001, minQ 20, minGQ 20, min-alleles 2, max-alleles 2" parameters. Finally, clean SNPs and indels were separated by vcftools.

#### 5.2 structural variation (SV) calling and genotyping.

Structural variants (SVs) were called using three classic WGS SV callers. In detail, The WGS data for each accession were mapped to reference using BWA-MEM with default parameter, and the duplications were marked using picard v2.25.7-0 with option "REMOVE_DUPLICATES= false". The sorted bam file was used for SV calling with Delly v0.8.7 (Rausch et al., 2012), smoove v0.2.8, and Manta v1.6.0 (Chen et al., 2016), respectively.

Delly identified deletions (DEL), insertions (INS), duplications (DUP), inversions (INV), and translocations (BND) for each individual by integrating the strategies of read depth, read pair, and split read for SV identification. SV were first called in each individual with parameter "-q 3 -r 20 -s 9 -x", gap regions greater than 100 bp were excluded and records with breakpoint offset less than 1kb and greater than 80% overlap rate with each other were merged. Single-individual SV genotyping was performed using SVs files merged in the previous step and their bam data. The final delly SVs calls file was generated by merging each genotyped individual with Bcftools v1.16 (Danecek et al., 2021) and filtering with the Delly "filter" option.

smoove wraps existing software (lumpy, svtyper, etc.) and adds internal read filtering to simplify calling and genotyping structural variants. SV calls were performed using the official Smoove pipeline with default options. SV calls were performed in four steps: (1) SV call from a single individual, (2) combine all site information, (3) combine all site information with reads quality to genotype SV called from a single individual, and (4) merges all the single sample VCF files to get the final file that contains all the variations. The same exclusion area as the Delly pipeline was used.

For the Manta pipeline, single-sample SV calls were performed using the officially recommended steps with default parameters, and the same regions as the two WGS SV pipelines mentioned earlier were excluded from the following analysis. The SURVIVOR v1.0.7 (Jeffares et al., 2017) method was used to combine the populations with the same SV type and breakpoints within 500 bp, and the minimum number of alleles involved in the recording was set to three (**Supplementary Tab.9**).

#### 5.3 Merging of all genotyping and filtering

To obtain high-quality full genotyping, a strict filtering pipeline was implemented. For Delly, we only SV sites with "PASS" tags were retained, while for smoove, we only kept sites that were not "IMPRECISE". For Manta, sites supported by fewer than two samples were removed. SURVIVOR v1.0.7 was used to combine SVs supported by at least two methods, and adjacent SVs were combined as a single SV if the distance between the start coordinate of one SV and the end coordinate of the other SV was less than 500 bp. Meanwhile, the SVs obtained by the different tools do not need to agree on the SV-type or strand, so that we can capture as many high-quality potential breakpoints as possible. Finally, the shared breakpoints of SV were screened, and SV sites with split-read (SR) values greater than three were retained.

##### 5.4 Evaluation of SV genotype data

To evaluate the sensitivity and accuracy of our structural variation (SV) genotype dataset, we incorporated five PacBio long-reads into our analysis. Briefly, comparing the contribution of long-reads and WGS data to the SV genotype data in an individual detected the sensitivity of the data, and assessing the accuracy depended on detecting the proportion of SV in WGS genotype SV data which could be remapped in long-reads data calls. Due to the limitations of short-read data, we evaluated only deletion (DEL) type SVs in this evaluation.

SV calls for long-reads data were performed using Sniffles v2.0.7 (Sedlazeck et al., 2018) and SVIM v1.4.2 (Heller & Vingron, 2019), because they have better accuracy and sensitivity (De Coster et al., 2019). First, we performed SV detection on each sample using the tools described above. To match the SV information of the WGS data, SVs with five minimum supported reads were retained. SURVIVOR v1.0.7 was used for merging, and the maximum allowed pairwise distance between breakpoints was 500 bp. SVs obtained from different tools do not have to agree on SV types or chains so that we can capture as many potential breakpoints as possible.

Based on the grouping information, we obtained the SVs data (WGS data called) that belonged to the same group as the long-read data. As the long-reads data for samples HB and JX were from the central region, we collected SV data only from samples from the same province. Finally, we graded the SV sets based on the degree of sharing among different groups and mapped them to the corresponding long-read SV data sets for this evaluation **(Supplementary Tab.10)**.

#### 6 SV distribution and annotation

To measure the distribution characteristics of SV in the reference genome, we examined overlapped region of the SV, and SVs with long below 30bp or higher than 100kb were excluded from the following analysis.

##### 6.1 Genomic background model

To assess the enrichment of genomic elements that overlapped with SVs, we generated 1000 randomly shuffled sets of SVs in non-gap regions of the reference genome (HB), as previously described (Quan et al., 2021; Sudmant et al., 2015). Briefly, we retained the SV type and length while shuffling the SVs present for each chromosome separately using BEDTools v2.30.0 (Quinlan & Hall, 2010), without intersection with the original SV position. These shuffled sets were used as *A. cerana* random background models to understand the tendency of SVs overlapped with genomic elements with null distribution and real. The Log2 fold change (LFC) method was used to measure enrichment, and we calculated the empirical p-value, which was considered significant if it was less than 0.05 after Bonferroni correction. Similarly, employing the same method, we generated a random TE background distribution to evaluate the bias of TE.

##### 6.2 SV distribution analysis

We investigated the presence of repeat elements in close proximity to structural variations (SVs) by analyzing the sequences flanking the SV breakpoint within a range of (+/- 100) base pairs. The intersections were identified using BEDTools v2.30.0, and the breakpoints were annotated based on their association with the overlapping repeat elements. The repeated elements were classified into six distinct groups, including low complexity repeats, simple repeats, non-LTR retrotransposon, long terminal repeat retrotransposon (LTR), DNA repeats (including MITE), and other repeats. Permutation tests were subsequently performed for each repeat class using the random background model. In this work, we do not consider translocations (BND) types of SV (**Supplementary Tab.9**).

##### 6.3 SV formation mechanisms

Previous reports have indicated that breakpoint junction sequences can be used to infer the formation mechanism of SV (Audano et al., 2019; Lam et al., 2010; Qin et al., 2021; Quan et al., 2021). We used a similar simplified algorithm to classify SVs. First, SVs with more than 50% overlapping with tandem repeats, simple repeats, or low complexity repeats were considered as expansion or contraction of VNTRs. Then, SVs were classified as NAHR if the 200bp sequence flanking the breakpoint was more than 80% identical, implying that the sequence flanking the breakpoint has homology. Finally, we defined SVs sites overlapping transposable element (TE) regions as TE-mediated mechanisms SV (see 6.2 methods). Based on the type of overlapping TE, they were divided into single transposable element (STE) and multiple transposable element (MTE).

##### 6.4 SNP LD ranking analyses of indels and SVs

To analyze the association of SVs with indels or SNPs, we performed linkage disequilibrium (LD) analysis and ranked the variants, following the methods used in previous studies (Gui et al., 2022; Stuart et al., 2016; Yang et al., 2019). After filtering (MAF>5%), we select the 150 closest SNP or indels sites upstream and downstream of the SV breakpoint, respectively. A total of 301 variants (the target variant is counted as a variant) are used to calculate the pairwise genotype LD (r2 value) using vcftools v0.1.16 with the "-- geno-r2" parameter. All pairwise LD values for each site were sorted in descending order, and the median value was selected as the threshold value above which was considered associated with the target variant. Target variants with less than one-third of the total ranks were classified as low-LD levels, and those with more than two-thirds ranks were classified as high LD levels. Variants between the two groups were classified as medium LD levels. In this study, only medium and high levels of LD-linkage levels were considered effective linkage.

#### 7 Population structure and phylogenies analysis

Analyses of population structure were performed using both SNP and SV datasets. To avoid the potential impact of LD on population structure and mixing analyses (Linck & Battey, 2019), we applied LD pruning and removed sites with minor allele frequency (MAF) below 0.05 (for SV data, we removed categories classified as Single). To ensure data integrity, the missing genotypes were imputed using Beagle v5.1(Browning et al., 2018) with hidden Markov model.

The program ADMIXTURE v1.3.1 (Alexander & Lange, 2011) was used to analyze the ancestry proportion and population structure of 525 *A. cerana* samples. We conducted 15 separate experiments with K ranging from 2 to 15 for each dataset and used a ten-fold cross-validation procedure to estimate the best K value. Principal component analysis (PCA) was performed using plink v1.90 (Purcell et al.,

2007) software to detect genetic relatedness and clustering patterns among individuals. The phylogenetic tree was constructed using IQ-TREE (Nguyen et al., 2015) program based on the SNP and SV data sets with 1000 bootstraps according to a maximum-likelihood method, respectively. We also visualized the SV dataset using the R package NetView 1.1 (Steinig et al., 2016) based on the IBS distance matrix generated by plink software with the "--distance" parameter. This package uses minimum spanning trees and mutual K-nearest neighbor graphs to generate networks. Closely related individuals form clusters that show distinct classes that are equivalent to genetically distinct populations or lineages in the population structure. The "Infomap" and "greedy" algorithms were used to detect clusters and combined with the ancestry fraction generated by ADMIXTURE to show the Ancestry of *A. cerana*.

#### 8 Population-stratified SVs and candidate adaptive genes

##### 8.1 stratified analysis

Using common SV genotype data (single SVs were removed), we applied the Weir and Cockerham estimator for *Fst* (Yang, 1998) based on VCFtools v0.1.16 to identify SVs stratified between peripheral and central groups **(Supplementary Tab.12)**. SV sites with concurrent in the top 1% of the *Fst* values were labeled as stratified SVs. To identify all candidate stratified genes that may be regulated by these SVs, we consider genes supported by two types of evidence for linear overlap with SV or the presence of SV within 2kb upstream of the gene. ML belongs to Sundaland *A. cerana*, was excluded in subsequent analyses.

##### 8.2 Genome-wide selective sweep analysis

Through two strategies, we identified SV loci with selective signals by examining nearby SNPs. To investigate independent habitat adaptation, we combined two ancestral lineages, CT-N and CT-S, which were distributed over most of the region, into a central group to rule out genetic drift **(Supplementary Fig.8 C)**. For the population differentiation strategy, the SNP nucleotide diversity ($\pi$) of multiple peripheral groups and single central groups was calculated using vcftools v0.1.16. The nucleotide diversity ratio was used to identify candidate peripheral local adaptation regions. For the composite likelihood ratio strategy, SweeD v4.0.0 (Pavlidis et al., 2013) was used to screen the selected region with a 1Kbp grid size. The composite likelihood ratio analysis can calculate within-group selection status, so it is necessary to split the SNP data set according to the grouping label. The regions corresponding to the top 5% likelihood value and $\pi$ ratio were selected, and the overlapping regions were identified as highly credible selected regions in the peripheral. Finally, we counted the distribution of SV in the selected regions **(Supplementary Tab.12)**.

#### 9 Outlier detection and cross-validation genetic-environmental association analysis

##### 9.1 Collection of bioclimatic variables

To infer the possible distribution range of *A. cerana* in different climatic periods, WorldClim Bioclim (v2.0) was downloaded as a predictor of the niche model (Fick & Hijmans, 2017). Based on the latitude and longitude of each bee sample, we extracted 19 temperature and precipitation related bioclimate variables (Bio1-bio19). These variables included: Annual Mean Temperature (Bio1), Mean Diurnal Range (Bio2), Isothermality (Bio3), Temperature Seasonality (Bio4), Max Temperature of Warmest Month (Bio5), Min Temperature of Coldest Month (Bio6), Temperature Annual Range (Bio7), Mean Temperature of Wettest Quarter (Bio8), Mean Temperature of Driest Quarter (Bio9), Mean Temperature of Warmest Quarter (Bio10), Mean Temperature of Coldest Quarter (Bio11), Annual Precipitation (Bio12), Precipitation of Wettest Month (Bio13), Precipitation of Driest Month (Bio14), Precipitation Seasonality (Bio15), Precipitation of Wettest Quarter (Bio16), Precipitation of Driest Quarter (Bio17), Precipitation of Warmest Quarter (Bio18), as well as elevation Precipitation of Coldest Quarter (Bio19).

#### 9.2 genetic-environmental association analysis

To identify top outliers within species as candidates for local adaptation, we applied a multi-model cross-validation approach based on previous studies (Heraghty et al., 2022; Jackson et al., 2020). First, we performed an environmental association analysis using a latent factor mixture model (LFMM2 in the LEA v3.0.0 package)(Gain & Francois, 2021), which employed the least-squares method to identify loci significantly associated with environmental variables. The sMNF was used to calculate population clusters (k) (Frichot et al., 2013) allowed us to refine different levels of stringency for analyses. After correction, significant environmentally relevant loci at a threshold of p.adjust ≤0.05 were considered candidates for local adaptive relevance.

Secondly, we used Bayenv2 as a cross-value outlier to validate the mechanism of the candidates from the LFMM. Different from LFMM, Bayenv2 uses population allele counts and controls for population structure using a genetic similarity correlation matrix based on the XTX statistic (similar to *Fst*) (Gunther & Coop, 2013). In Bayenv2, the Bayesian factor (BF) was provided for each locus as evidence in support of the model with the environmental parameter added over the neutral model. We also evaluated the transformed rank statistic (ρ) as one of the evaluation criteria. According to the recommendations of the Bayenv2 (https://bitbucket.org/tguenther/bayenv2_public/src/master/), variant sites with high BFs or ρ far away from zero are considered robust. In our data, the threshold of 5% BF and 10% |ρ| were used for the acquisition of candidate loci **(Supplementary Fig.11; Supplementary Tab.13)**.

Candidate sites marked as significant in both LFMM and Bayenv2 were used for further analysis. Here, we only consider the sites where genes exist within 2 kb near the SV breakpoint. The initial list of 95 GO terms ($P < 0.01$, including 57 for temperature and 38 for precipitation) was obtained and subsequently reduced to 50 biological terms using the Revigo Abstract Web tool (26 for temperature-related and 26 for precipitation-related terms; **Supplementary Tab.14**)(Supek et al., 2011), certain outlier genes were excluded from the GO analysis due to a lack of annotation(Walsh et al., 2022) (**Supplementary Tab.14**).

#### 10. Functional exploration of a 330 bp deletion in an environment-related gene

##### 10.1 SV validation

To validate the structural variants (SVs), we performed a polymerase chain reaction (PCR) assay. First, we retrieved the genome deletion carrying sample from AB group. The head and thorax were preserved in liquid nitrogen after the removal of the intestine, and the abdomen was retained for DNA extraction.

A 500-bp genomic sequence around the DEL00042633 breakpoint was extracted from the HB assembly. Then, Primer6 (v.0.4.0) was used to calculate primer pairs flanking the breakpoints in these regions for PCR amplification (**Supplementary Fig.15**).

#### 10.2 Effects of genomic deletions on gene function

Based on the PCR validation results, six bee samples (including three "deletion" and three wild types) were subjected to RNA sequencing on the Illumina high-throughput sequencing platform NovaSeq 6000, where each sample was separately sequenced according to the head and chest regions. Gene expression was calculated using FeatureCounts and DESeq2 as described in section 4.3.

### 11 Visualizations

The circos plot of the *A. cerana* genome was drawn using the TBTOOLS package (Chen et al., 2020). The basic statistic plots such as the histograms, boxplots, densities, lines, and dots were drawn using R/ggplot2 (Wickham, 2011), and the basic drawing of the violin diagram relies on the R/ggpubr (https://github.com/kassambara/ggpubr). Heat maps were generated using R/pheatmap (https://github.com/raivokolde/pheatmap) and genome visualization was performed using IGV software (Robinson et al., 2022). GO enrichment and KEGG analysis of *A. cerana* were performed using "enrichGO" and "enrichKEGG" functions of R package clusterProfiler (Yu et al., 2012).
